# A collicular map for touch-guided tongue control

**DOI:** 10.1101/2024.04.08.587629

**Authors:** Brendan S. Ito, Yongjie Gao, Brian Kardon, Jesse H. Goldberg

## Abstract

Accurate goal-directed behavior requires the sense of touch to be integrated with information about body position and ongoing motion^1,2,3^. Behaviors like chewing, swallowing and speech critically depend on precise tactile events on a rapidly moving tongue^4,5^, but neural circuits for dynamic touch-guided tongue control are unknown. Using high speed videography, we examined 3D lingual kinematics as mice drank from a water spout that unexpectedly changed position during licking, requiring re-aiming in response to subtle contact events on the left, center or right surface of the tongue. Mice integrated information about both precise touch events and tongue position to re-aim ensuing licks. Surprisingly, touch-guided re-aiming was unaffected by photoinactivation of tongue sensory, premotor and motor cortices, but was impaired by photoinactivation of the lateral superior colliculus (latSC). Electrophysiological recordings identified latSC neurons with mechanosensory receptive fields for precise touch events that were anchored in tongue-centered, head-centered or conjunctive reference frames. Notably, latSC neurons also encoded tongue position before contact, information important for tongue-to-head based coordinate transformations underlying accurate touch-guided aiming. Viral tracing revealed tongue sensory inputs to the latSC from the lingual trigeminal nucleus, and optical microstimulation in the latSC revealed a topographic map for aiming licks. These findings demonstrate for the first time that touch-guided tongue control relies on a collicular mechanosensorimotor map, analogous to collicular visuomotor maps associated with visually-guided orienting across many species.

## Introduction

If your pinky nicks your coffee cup as you extend your arm to reach for it, you immediately reposition your fingers laterally to optimally grasp and lift the cup. Similarly, as you eat and drink, tactile feedback enables your tongue to dexterously handle food and water for chewing and swallowing while maintaining airway patency and preventing tongue injury. Impairments in the tongue’s ability to handle food and water result in aspiration pneumonia, a leading cause of death in neurological disease^6,7^. Just as fingertip anesthesia impairs grasping even with visual feedback intact^8,9^, blockade of tactile feedback to the tongue impairs chewing, swallowing and speech^4^. Yet the neural circuits underlying touch-guided tongue control remain unknown.

### Tactile events guide ongoing tongue movements

To test if mice use tactile feedback for tongue control, we developed a touch-guided lick task that required mice to re-aim licks in response to subtle contact events on the tongue surface. We first trained head-fixed mice to withhold licking for 1 s to receive an auditory cue, and to contact a water spout aligned to the center of the facial midline within 1.3 s to earn a water reward. Mice naturally lick in rhythmic bouts of 4-8 licks at ∼8 Hz^10^. We detected spout contacts in real time and randomly displaced the spout to the left or right by approximately one half of the tongue width (1.1 mm on average) between the first (L1) and second (L2) licks of a bout (Fig. 1a, ∼400 trials per session; spout moved to left, center, or right positions with equal probability, Methods). Importantly, the subtle spout displacements between L1 and L2 did not cause spout misses on L2, but rather evoked unexpected nicks at distinct locations on the tongue surface (Fig. 1b-d).

**Figure 1.**
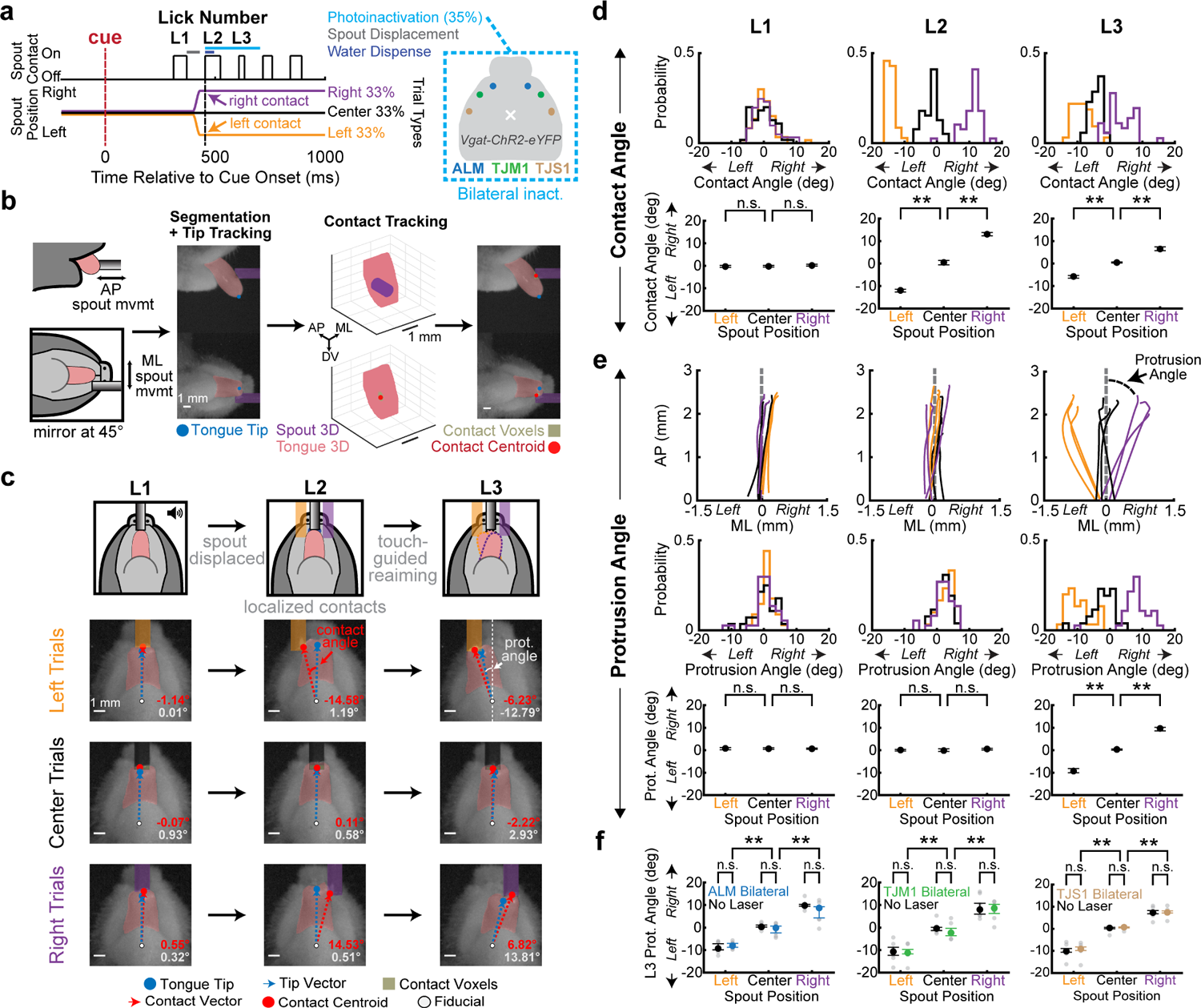
Lingual cortical areas are not required for touch-guided re-aiming. **a**, Left, Trial structure. An initially centered water spout was randomly moved to center, left or right positions between the first (L1) and second licks (L2) of a bout. On 35% of trials, L2 contact triggers laser-on for 250 ms. Right, Optical fibers were implanted bilaterally over ALM, TJM1 or TJS1 in *Vgat-ChR2-eYFP* mice. **b**, Left, Schematic and still frame of L2 contact on a spout-right (purple) trial. Side and bottom views were filmed at 1 kHz frame rate, two convolutional neural networks segmented the tongue (pink mask), and tongue tip location (blue dot) was computed from the 3-D hull tongue reconstruction on each frame. Right, Contact centroid (red dot) was defined as the centroid of all contact voxels (brown mask). **c**, Bottom view still frames at L1-L3 contact onset across trial types. Contact site was defined as the angle between the tip vector (blue dashed line, fiducial to tongue tip) and contact vector (red dashed line, fiducial to contact centroid). Protrusion angle was defined as the angle between the tip vector and facial midline. **d**, Top, L1-L3 contact angle distributions on left (gold), center (black) and right (purple) trials for a single session. Bottom, L1-L3 contact angles across mice (*n* = 14 mice). Note the impact of spout position on L2 contact angles. **e**, Top, Example L1-L3 tongue tip protrusion trajectories. Middle and Bottom, Same as **d**, except for L1-L3 protrusion angles. Note the impact of spout position on L3 protrusion angles. **f**, L3 re-aiming was intact during photoinactivation of either ALM, TJM1 and TJS1. L3 protrusion angles for no laser (black) trials and ALM (blue, *n* = 7 mice), TJM1 (green, *n* = 6 mice) and TJS1 (brown, *n* = 6 mice) inactivation trials. Gray dots are medians of individual mice. Data in **d-f** are median ± IQR. **corrected *p* < 0.01, shuffle test; n.s., not significant. Exact statistics are in Supplementary Tables 1 and 3.

To precisely quantify both 3D tongue kinematics and nick sites, we combined dual-plane kilohertz frame-rate tongue imaging, deep-learning-based image segmentation, and visual hull reconstruction to estimate the tongue 3D volume and tip position in each frame, resulting in millisecond-resolution tongue tip trajectories for each lick^11^ (Fig. 1b, Methods). Next, to determine the precise location on the tongue surface that contacted the spout on each lick, we embedded a 3D model of the spout in the same space as the 3D hull reconstruction of the tongue, and identified the centroid of the reconstructed tongue voxels that neighbored or overlapped with spout voxels (Fig. 1b, Methods). We defined a fiducial point at the base of the jaw that defined the midline of each animal, and then quantified the spout-tongue contact site on each lick as the angle between two vectors: from the fiducial point to the tongue tip (i.e., the tip vector) and the fiducial point to the contact centroid (i.e., the contact vector) (Fig. 1c). We chose an angular coordinate system because it captures tongue contact site variance in both the mediolateral and anteroposterior axes with a single term (Methods). Together, these methods allowed us to track both nick sites and lick kinematics with millisecond precision.

In all mice tested, left or right nicks on L2 (Fig. 1d) caused immediate re-aiming of the next lick (L3, Fig. 1e, Supplementary Video 1, Methods), suggesting that mice naturally use touch to guide tongue aiming within lick bouts. Touch-guided re-aiming did not depend on water dispensation at contact (Extended Data Fig. 1), and the rhythm of the lick bout was not affected by the process of touch-guided re-aiming (Extended Data Fig. 2a), showing that the sensorimotor process linking tactile feedback circuits to premotor tongue steering circuits occurs rapidly, within a single inter-lick interval (median latency from L2 contact onset to L3 protrusion onset: 141 ms [IQR of 127-148 ms], *n* = 14 mice).

### Orofacial sensory and motor cortices are not required for touch-guided re-aiming

Touch-guided grasping and visually-guided reaching depend on sensory and motor cortical processing^12–14^. Three sensorimotor lingual cortical areas, tongue-jaw sensory cortex (TJS1), tongue-jaw motor cortex (TJM1) and anterolateral motor cortex (ALM) are implicated in diverse lick aiming tasks^11,15–18^. To test if touch-guided tongue aiming similarly requires cortical pathways, we used VGAT-ChR2-EYFP mice to bilaterally photoinhibit TJS1, TJM1 or ALM^11,15–17^. Photoinhibition for each region was initiated at the moment of L2 spout contact on 35% of trials and lasted for 250 ms, ensuring inhibition from L2 contact through L3 re-aiming (Fig. 1a, Extended Data Fig. 3, Methods). Licks produced during ALM and TJM1 inactivation exhibited reduced durations and pathlengths (Extended Data Fig. 2e-j), consistent with past reports^11,16^, suggesting that photoinhibition effectively impaired the function of the relevant brain regions. Yet surprisingly, touch-guided re-aiming on L3 was completely intact both during photoinactivation of TJS1, TJM1, and ALM (Fig. 1f, Supplementary Video 2) and after complete bilateral lesion of all three areas at once (Extended Data Fig. 4, Supplementary Video 3), suggesting that lingual cortical areas are not required for touch-guided tongue aiming.

### The latSC is important for touch-guided re-aiming

Across vertebrates, the superior colliculus (SC) is involved in stimulus-guided orienting^19–22^. The SC receives direct visual input from the retina^20,23^ and contains a topographically-organized visuomotor map in which each region of the visual field maps to a corresponding territory in the contralateral SC^24,25^. We hypothesized that a lateral region of the superior colliculus (latSC) recently associated with directional licking^26–29^ may be analogously important for touch-guided tongue control.

To test the role of the latSC in touch-guided tongue aiming, we used Vgat-cre mice to express channelrhodopsin-2 (ChR2) in GABAergic inputs to the latSC from the substantia nigra pars reticulata (SNr) and implanted optic fibers over the latSC. This approach was shown to inhibit latSC discharge and impair licking in past work^26^. We bilaterally photoinhibited the latSC by photoactivating SNr terminals in the latSC from L2 contact onset through the entire duration of L3 on 35% of trials (Fig. 1a & 2a, Methods). In contrast to inactivation of ALM, TJS1 or TJM1, bilateral latSC inactivation immediately interrupted lick bouts (Fig. 2b, Extended Data Fig. 2b-d, Supplementary Video 4), consistent with prior work^26^.

**Figure 2.**
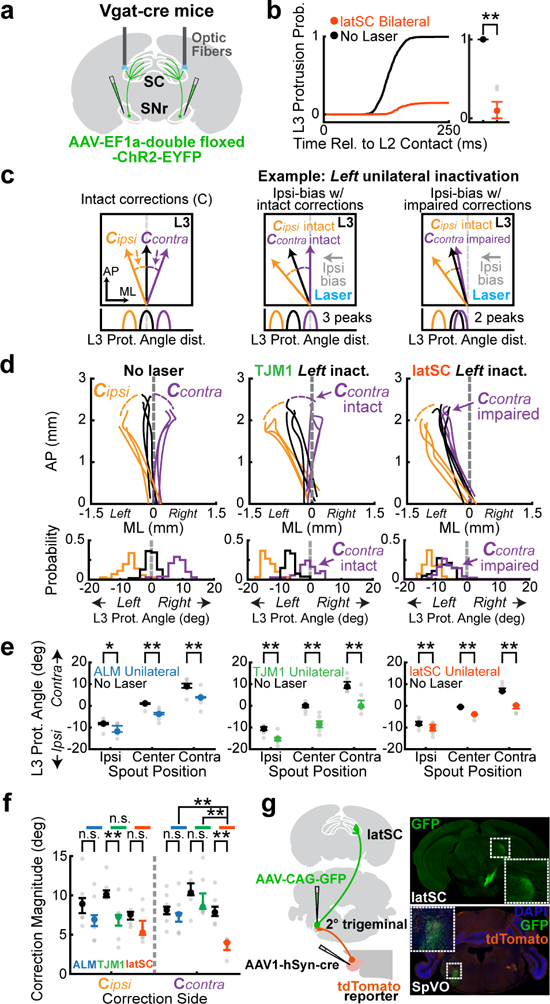
Photoinactivation of the lateral superior colliculus (latSC) impairs touch-guided re-aiming. **a**, latSC photoinhibition strategy. Photoinhibition timing executed as in Fig. 1a. **b**, Left, Cumulative L3 protrusion probability for no-laser (black) and latSC-inactivated trials (red-orange) across mice. Right, L3 protrusion probability (*n* = 6 mice). **c**, Schematics showing three hypothetical ways left unilateral photoinhibition at L2 contact could plausibly affect L3 protrusion angles. Unilateral activations could have no effect (left), could cause an ipsilateral steering bias across all trial types but preserve corrections (center), or could cause both an ipsilateral bias and impair contralateral corrections on L3 (right). Contralateral (C_contra_) and ipsilateral (C_ipsi_) corrections on L3 were defined as the difference in median protrusion angle between the center and contralateral, or ipsilateral, trials for each mouse. **d**, Example L3 protrusion trajectories (top) and protrusion angle distributions (bottom) from single sessions for no laser (left) trials, and left TJM1 (center) or latSC inactivation (right) trials. Note that C_contra_ and C_ipsi_ were preserved with left TJM1 inactivation, whereas left latSC inactivation specifically impaired C_contra_. **e**, L3 protrusion angles for laser-off (black) and laser-on trials; unilateral ALM (blue, *n* = 7), TJM1 (green, *n* = 6), and latSC (red-orange, *n* = 6). Note all inactivations caused an ipsilateral bias. **f**, C_contra_ and C_ipsi_ re-aiming magnitudes for ALM, TJM1 and latSC photoinactivations. Colors are as in **e.** Note that latSC photoinactivation specifically impaired C_contra_. **g**, Left, Tracing schematic. Right, Axonal projections from SpVO to the latSC (top, green) and axonal projections from 1° trigeminal to SpVO (bottom, orange) with GFP-labelled SpVO cell bodies (green). Data in **b**, **e** and **f** are median +/- IQR. Gray dots in **e-f** are medians of individual mice. *corrected *p* < 0.05, **corrected *p* < 0.01, shuffle test; n.s., not significant. Exact statistics are in Supplementary Tables 4, 7, and 8.

We reasoned that unilateral inactivation experiments may help clarify the precise role of the latSC in touch-guided re-aiming. In our task, tactile signals from L2 nicks must somehow reach tongue steering circuits before L3. Any interruption of this sensorimotor path may block touch-guided re-aiming on L3. Unilateral inactivations of multiple brain regions, including TJM1, ALM or latSC, are all known to cause an ipsilateral steering bias^17,27,30,31^. Thus, if any one of these brain regions can contribute to lick aiming but is not necessary for reacting to the touch event, then unilateral inactivation of that region during and after the L2 nick should bias all L3 protrusions ipsilaterally, without eliminating re-aiming due to the L2 nick event. So, over a session of left, center and right spout contacts on L2 with unilateral photoinactivation in those regions, three distinct peaks in the L3 protrusion angle distributions would still exist on both intact and unilateral inactivation trials, but the peaks with unilateral inactivation would all be biased ipsilaterally towards the inactivated side (Fig. 2c, compare left and middle panels). This outcome would suggest that while the inactivated brain region can contribute to tongue steering, a distinct circuit is driving the touch-guided re-aiming process. On the other hand, if a brain region is an essential part of the touch-to-steering circuit, then unilateral inactivation of that region might block the capacity to re-aim in response to contralateral, but not ipsilateral, nicks^27,32^. As a result, the peak representing contralateral trials would largely merge with the peak for center trials, leaving only two peaks (Fig. 2c, compare left and right panels). This result would also suggest that other steering circuits may not be sufficient to implement contralateral touch-guided corrections on their own.

We performed unilateral photoinactivation of ALM, TJM1 or the latSC in separate batches of mice with the same duration and timing as bilateral photoinactivation to test these hypotheses. We defined the correction magnitude for each animal as the difference between the median L3 protrusion angle for the center condition and the median L3 protrusion angle for either the contralateral or ipsilateral condition, which were defined relative to the inactivated hemisphere to allow comparisons. Unilateral ALM, TJM1 and latSC inactivations minimally impaired L3 lick kinematics (Extended Data Fig. 5), and as expected, biased licks ipsilaterally^30,32–34^ (Fig. 2d-e, Supplementary Video 5). But critically, unilateral latSC, but not ALM or TJM1, inactivation selectively impaired contralateral re-aiming, while preserving ipsilateral re-aiming (Fig. 2d-f, Supplementary Video 6, Supplementary Table 8, 4.0° [IQR of 2.2-4.5°] median magnitude of contralateral correction with unilateral latSC inactivation, 7.9° [IQR of 6.8-9.3°] control). Notably, this result resembles impairments from unilateral SC inactivations in saccade tasks, where orienting to contralateral visual targets is specifically impaired^33,34^. These findings suggest that the latSC is part of the neural pathway that links tactile information from L2 to re-aiming commands for L3.

### An afferent pathway from the tongue to the latSC

If the latSC is involved in processing touch signals from the tongue to guide next lick aiming, then the latSC should receive input from tongue sensory afferents. To test this idea, we carried out viral tracing experiments with tongue injections. Oral sensory neurons reside in the primary (1°) trigeminal ganglion and project to secondary (2°) trigeminal nuclei in the brainstem^35^. To test if the latSC receives input from 2° trigeminal nuclei that, in turn, receives input from 1° tongue trigeminal neurons, we injected a Cre-expressing adeno-associated virus (AAV) into the tongues of *Rosa26^LSL-tdTomato^*mice to fluorescently label 1° trigeminal neurons innervating the tongue. We identified axon terminals in the dorsal subdivision of spinal trigeminal nucleus oralis (SpVO), a 2° trigeminal region known to project to the hypoglossal nucleus and the latSC, suggesting a role in touch-guided tongue control^36,37^. We then injected a GFP-expressing AAV into this part of the SpVO and identified projections to the contralateral latSC (Fig. 2g, n = 3 mice). The discovery that the latSC receives input from a lingual territory of SpVO raises the possibility that the latSC might process tactile feedback to guide ongoing licking.

### The latSC encodes the location of contact events on the tongue

To test for possible roles of the latSC in processing tactile feedback, we recorded latSC neural activity during the touch-guided lick task (*n* = 553 neurons, 64 sessions, 6 mice). Many latSC neurons exhibited touch-site-dependent (left/center/right) changes in activity in the first 100 ms after L2 contact (Fig. 3a-b, Extended Data Fig. 6a-b, *n* = 279/553 neurons, Methods), with most touch-sensitive neurons selective for contralateral contacts (Fig. 3b-c, Extended Data Fig. 6a-b, *n* = 184/279 neurons with significant contralateral selectivity; Methods). The relative proportion of neurons with contralateral contact selectivity was significantly higher at posterior versus anterior latSC recording locations (Extended Data Fig. 6c, Supplementary Table 12), suggesting selectivity is graded along the anteroposterior axis in the latSC. Interestingly, a similar anteroposterior gradient for nasal to temporal visual selectivity is observed in the SC of many species^19,2019,21,38^.

**Figure 3.**
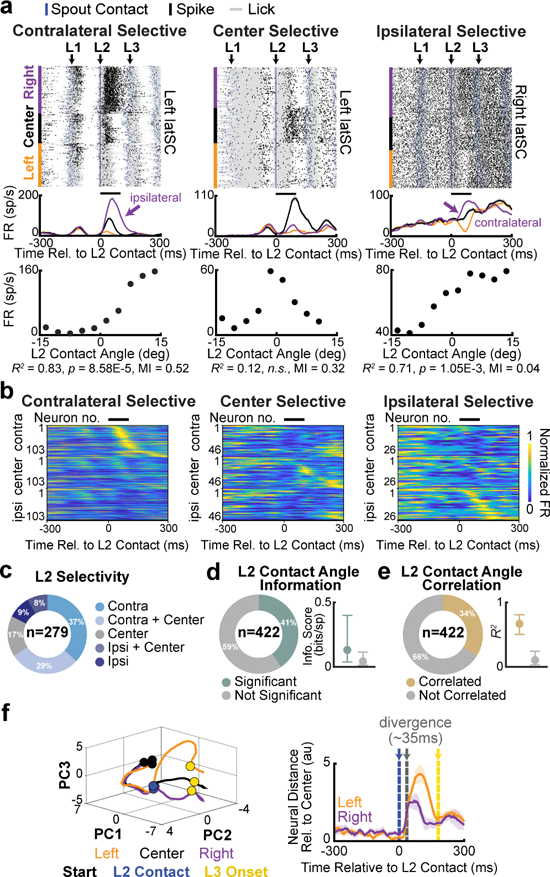
latSC neural activity encodes the location of contact events on the tongue surface. **a**, Spike rasters (top row) and rate histograms (middle row) from example latSC neurons selective for contralateral (left), center (middle) and ipsilateral (right) L2 contacts. Data are aligned to L2 contact onset and sorted by condition, then by L2 contact angle within each condition. Blue ticks and gray lines in each row of the raster signify spout contact and lick duration, respectively. Colors as in Fig. 1a. Black bar above the histogram indicates the spike window (100 ms following L2 contact) used to assess differences in firing rate between conditions. Bottom row, relationship between L2 contact angle (binned in 3°) and the mean firing rate of each neuron for each bin in the spike window. **b**, Normalized mean firing rates of latSC neurons selective for contralateral (left, *n* = 103/279), center (middle, *n* = 46/279) or ipsilateral (right, *n* = 26/279) L2 contacts aligned to L2 contact. Each neuron was normalized to its peak firing rate across all 600 ms, but sorting order across conditions was determined by peak activity in the spike window only in the condition those neurons were selective for. **c**, Percentage of L2 contact-modulated neurons selective for contralateral (cyan), contralateral and center (light blue), center (gray), ipsilateral (dark blue), and ipsilateral and center (blue) contacts. **d**, Left, Percentage of L2 contact-modulated neurons with significant contact angle information (green). Right, Median ± IQR contact angle information in bits/spike for neurons with significant contact angle information. **e**, Left, Percentage of L2 contact-modulated neurons whose activity was correlated with L2 contact angle (yellow). Right, Median ± IQR *R^2^* across neurons whose activity was correlated with L2 contact angle. **f**, Left, Top three principal components (PCs) of trial-averaged latSC population activity for each condition. Black, blue, and yellow dots indicate trajectory onset, L2 contact onset, and L3 onset, respectively. Right, Euclidean distance between the neural trajectories for left (gold) and right (purple) trials relative to center trials averaged across mice. Shading indicates bootstrapped s.e.m. of the neural distances. Blue, gray, and yellow lines indicate L2 contact onset, divergence, and L3 onset, respectively.

Principal component analysis of latSC population activity showed that neural trajectories following left or right contacts significantly diverged from the neural trajectory following center contacts very rapidly (Fig. 3f, 35 ms after contact [IQR of 30-35 ms], Methods). Notably, contact response latencies could be faster in individual latSC neurons (mode of neuronal latency distribution: 16 ms, Extended Data Fig. 6d, Methods). Thus contact representations arose during L2 contact (median L2 contact duration: 61 ms [IQR of 47 - 73 ms]) and well before the initiation of L3 (median latency from L2 contact to L2 retraction offset: 69.5 ms [IQR of 49.5-80 ms]; median latency from L2 contact to L3 protrusion onset: 141 ms [IQR of 127-148 ms]).

We wondered if the precise location of contact events on the subregions of the tongue surface sampled in our task influenced latSC discharge in neurons responsive to L2 contact. We visualized tuning to contact locations by plotting the firing rate of single latSC neurons following L2 contact against L2 contact angle (Fig. 3a (bottom), Methods), defined as the angle between the tip vector and the contact vector at the moment of spout contact. We assessed the significance of receptive fields in two ways. First, we computed a mutual information metric that quantifies site-specific contact information present in spikes^39^. Using this metric, 41% of all neurons significantly encoded contact angle information (*n* = 171/422, Fig. 3d), with some neurons exhibiting highly selective responses (left and middle example in Fig. 3a). Second, we correlated latSC single unit firing rates with L2 contact angle (Fig. 3a (bottom)) and found that 34% (*n* = 143/422) of neurons that were modulated following L2 contact exhibited significant tuning, including many with strikingly strong linear fits (Fig. 3e, Methods). Together, these complementary analyses show for the first time that latSC neurons can exhibit precise tactile responses to tongue contacts and exhibit significant selectivity for contact angle. The diverse range of tuning widths for contact angle resembles the variation in receptive field sizes previously observed for visual and auditory stimuli in the rodent SC^25,40,41^.

To test the behavioral relevance of these fine scale tactile representations, we plotted the magnitude of L3 re-aiming against the precise contact angle on L2 and identified a remarkably strong correlation that was significant both within and across trial types (Extended Data Fig. 7). Larger L2 contact angles resulted in larger protrusion angle changes between L2 and L3 (Extended Data Fig. 7c-d), even for only slightly off-center nicks (Extended Data Fig. 7d), demonstrating that mice used the precise locations of nick sites on L2 to adjust their aim for L3.

### latSC neurons represent contact location in tongue-centric, head-centric and conjunctive coordinate reference frames

For tactile feedback to be transformed to an estimate of spout position for aiming the next lick, the mouse needs to know not just where on the tongue surface the contact occurred but also the position of the tongue at the moment of contact. For example a left nick during a straight lick specifies a spout to the left of the head, but the same left nick during a right-aimed lick specifies a centered spout. Touch based re-aiming may require a coordinate transformation from a tongue- to a head-based reference system.

To examine the coordinate systems for lingual contact representations, we created a new ‘recentering’ task that decouples the nick site from the spout position. In this task, we produced left/right spout displacements on L2 that were followed by recentering spout displacements between L3 and L4 on 35% of trials (Fig. 4a). Mice reliably used L4 contacts to re-aim L5, showing that nicks on laterally-aimed licks can cause re-aiming to center (Fig. 4b, Supplementary Video 7). Note that tongue- and head-based reference frames make distinct predictions for neural responses to L2 and L4 contacts in the recentering task. A tongue-centric frame would encode the position of the nick site in a coordinate system anchored to the tongue, which would predict similar nick site selectivity on L2 and L4, even for nicks specifying distinct spout locations relative to the head (Fig. 4c, top). In contrast, a head-centric frame would encode the position of the spout in space relative to the head, which would require tactile responses to be integrated with information about the position of the entire tongue at the moment of contact (Fig. 4d, top).

**Figure 4.**
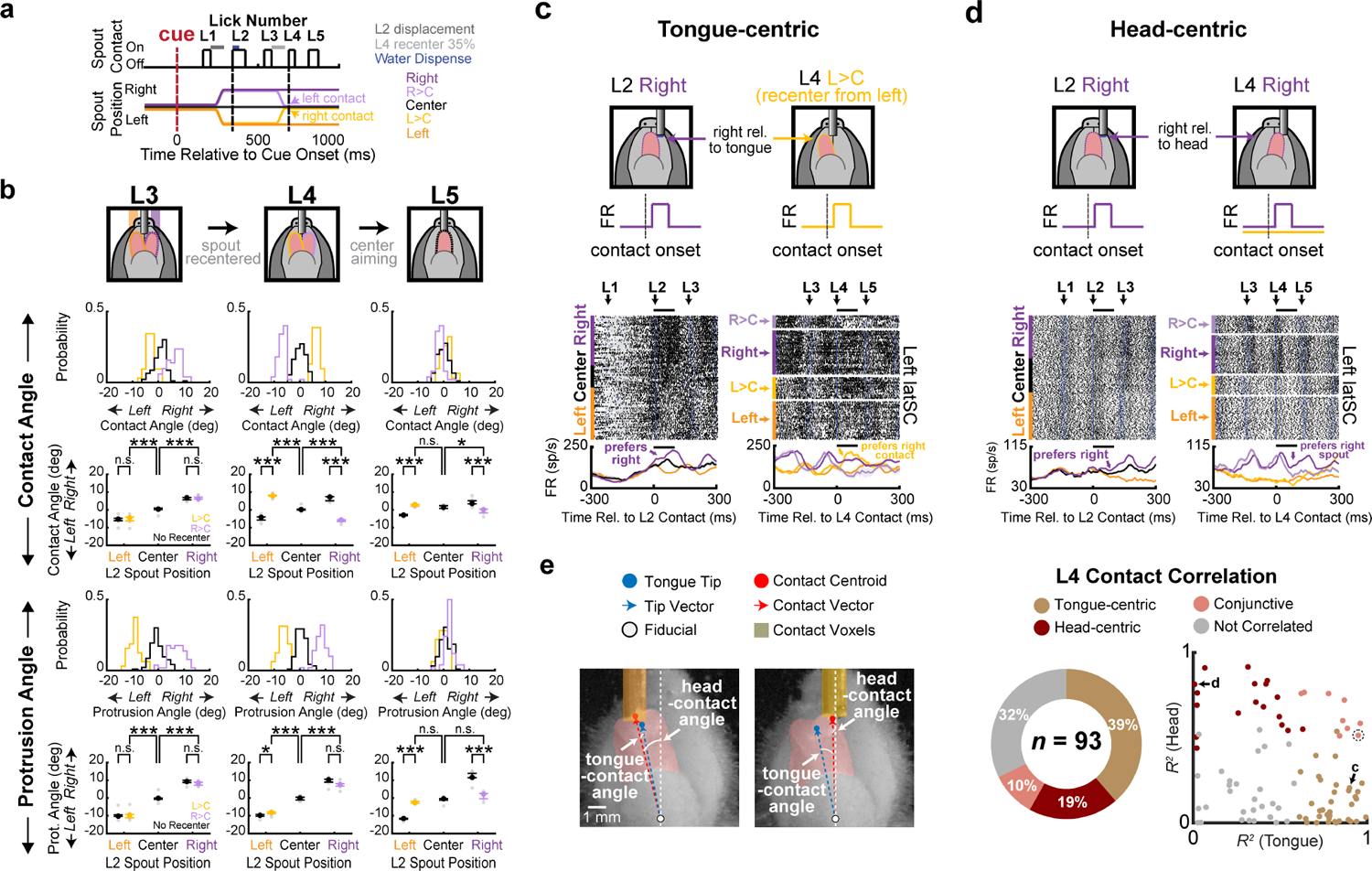
latSC neurons encode contact location in tongue-centric, head-centric and conjunctive coordinate frames. **a**, Trial structure in recentering sessions. The spout was randomly (35% of left/right trials) returned from left/right back to the center position between the third (L3) and fourth (L4) licks of a bout. Recentering sessions had five trial types: the spout could remain left (orange), center (black), or right (purple), or the spout could return from left to center (Left>Center/L>C, light orange) or right to center (Right>Center/R>C, light purple). Note that, L>C trials evoke right nicks on L4 and R>C trials evoke left nicks on L4. **b**, Top, Schematic of L3-L5 contacts and re-aiming on recentering trials. Middle, L3-L5 contact angle histograms (top) from an example session and median ± IQR across all mice (bottom, *n* = 5). Bottom, Same as in top, but for L3-L5 protrusion angles. Note that lateralized contacts on L4 lead to center re-aiming on L5. **c**, Top, Illustration of a tongue-centric neuron activated by the same contact location on L2 and L4. Bottom, Spike rasters and rate histograms for an example latSC neuron aligned to L2 (left) and L4 contact (right). Note that this neuron was activated by right-relative-to-tongue contacts (tongue-centric) on both L2 (right trials) and L4 (L>C trials). Colors as in **a**. **d,** Same as **c**, but for neurons activated by right-relative-to-head contacts on L2 and L4 (head-centric). **e,** Left, still frames showing examples of tongue-contact and head-contact angle. Colors as in Fig. 1b. Middle, Percentage of L2- and L4-modulated neurons significantly correlated with contact angle in a tongue-centric frame (gold), head-centric frame (red), both (pink), or neither (gray). Right, scatter plot of *R^2^* values from each neuron’s trial-by-trial correlations between mean firing rate following L4 contact and contact angles computed in tongue-centric (x-axis) or head-centric (y-axis) frames. The circled conjunctive neuron corresponds to the neuron in **Extended Data** Fig. 8g. Exact statistics are in Supplementary Tables 10 and 11.

To determine how contact representations depended on tongue position, we first noted that many neurons exhibited selectivity for contacts on specific licks in the bout (Extended Data Fig. 8a-f). We identified neurons during recentering sessions that exhibited nick site-selective responses to both L2 and L4 (Extended Data Fig. 8c; *n* = 93/359 neurons, *n* = 4 mice). Next, we compared L2 and L4 nick-site selectivities for each neuron and discovered that some exhibited tongue position-independent nick-site selectivity. For example, the neuron in Fig. 4c increased its firing rate following right surface nicks on both L2 and L4, even though the tongue was at distinct positions at the moments of contact. Yet the nick responses in other neurons depended on tongue position. For example, the neuron in Fig. 4d was activated by contacts on both L2 and L4 that specified a spout to the right of the head, but was not activated by right nicks on L4 that indicated a centered spout.

To quantify the contributions of both head- and tongue-based coordinate frames to each neuron’s response profile, we correlated each neuron’s L4 tactile response to contact angle encoded in tongue-(relative to tongue midline) or head-(relative to head midline) centered coordinates (Fig. 4e). The neural responses of some neurons were strongly anchored in a tongue-based reference frame, others in a head-based frame, and yet others exhibited conjunctive tuning to both tongue- and head-based coordinate systems (Extended Data Fig. 8g). Interestingly, circuits that implement coordinate transformations in other systems also exhibit neuronal representations anchored in both single and conjunctive reference frames^42–46^.

We next reasoned that the process of transforming tactile signals from a tongue-to-head based reference frame may impose a delay. We hypothesized that neurons with tongue-centered tactile responses might have lower contact latencies than neurons with head-centered responses, as head-centered neurons need to additionally weigh information about tongue position. Indeed, we found that neurons with a head-based reference frame had significantly longer contact latencies (Extended Data Fig. 8h), similar to past work examining analogous ego-to-allocentric sensorimotor transformations in the *Drosophila* central complex^47,48^.

Implementation of a tongue-to-head based coordinate transformation for tactile representations requires information about the position of the tongue, which could arise as a result of premotor signals, proprioceptive feedback and/or efference copy. One indicator of tongue position is phase in the lick cycle, which we quantified using the volume of the tongue visible outside of the mouth (Extended Data Fig. 9a-d, Methods). latSC neuronal activity strongly co-varied with lick phase (Extended Data Fig. 9a-c), as previously reported^26,27^ and neurons were locked to lick phase with a wide range of lags (Extended Data Fig. 9d). Yet to implement lateralized touch-guided corrections in our task, the latSC additionally needs detailed information about tongue position in the mediolateral plane, which is unspecified by lick phase alone. We used generalized linear models (GLMs) to test if fine details of 3D tongue position were encoded in the activity of individual latSC neurons in a brief interval preceding L4 contact (Extended Data Fig. 9e-f, 50 ms prior to L4 protrusion to L4 contact, Methods)^47,48^. Remarkably, L4 tongue position was strongly encoded in many latSC neurons prior to L4 contact (Extended Data Fig. 9g-i), and the mediolateral position of the tongue tip provided the strongest contribution to latSC spiking immediately prior to L4 contact on average (Extended Data Fig. 9h-i). Together, these analyses show that the latSC has the necessary information with the appropriate timing for implementing a tongue- to head-based coordinate transformation important for using the sense of touch to localize the spout in space, which in turn is important for aiming the next lick.

### A topographic map for tongue aiming in the latSC

To test if the latSC can play a causal role in lick generation and aiming, we optogenetically microstimulated four different regions spanning 800 μm along the AP axis of each latSC hemisphere for 400 ms (Fig. 5a, c (top), Extended Data Fig. 10a-c, Methods). Photostimulation of the latSC at each site reliably and rapidly elicited tongue protrusions and lick bouts even when photostimulation occurred outside of any task structure (Fig. 5b, e-f, Extended Data Fig. 10f-m). Critically, the precise photostimulation site was significantly correlated with the protrusion angle of the first stimulation-evoked lick (Fig. 5c-d, Extended Data Fig. 10d-e, Supplementary Video 8, Methods). Posterior stimulation sites directed the tongue more laterally and anterior sites more medially on the contralateral side (Fig. 5c-d, Extended Data Fig. 10d-e). A similar topographic layout for saccades and visually-guided orienting is observed in the SC across many species^19,2019,21,38^. Together with our electrophysiological data (Extended Data Fig. 6c), these findings show that the latSC contains a mechanosensorimotor map for touch-guided tongue control with a similar topography to visuomotor maps for saccades and orienting^19,20^.

**Figure 5.**
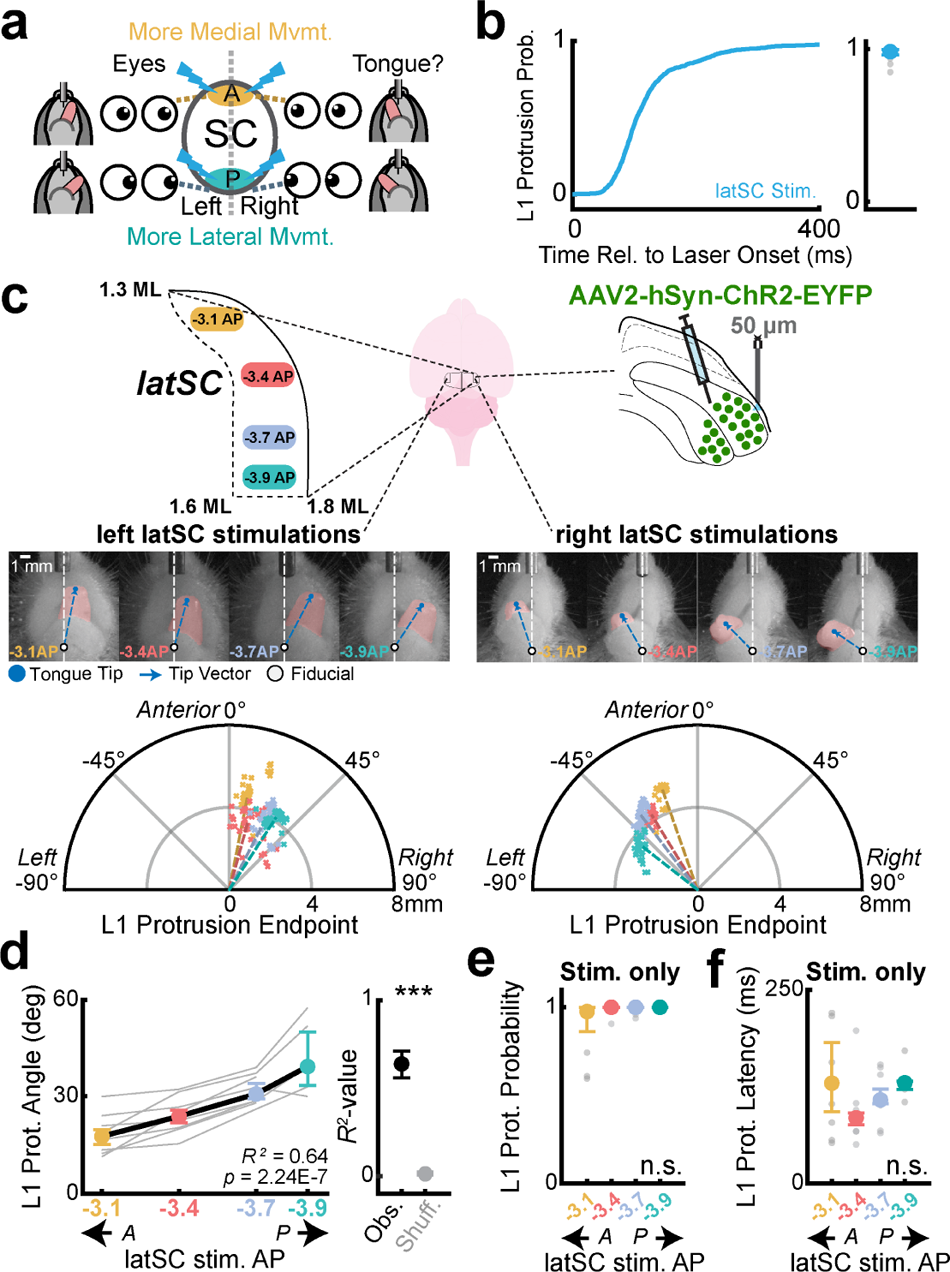
latSC photostimulation reveals a topographic map for tongue aiming. **a**, Hypothesis schematic. In many species, stimulation of posterior (P) sites in SC evoke more lateralized shifts in contralateral gaze than anterior sites (A). Multisite photostimulation in the latSC tested for similar functional tongue control. **b**, Left, Cumulative probability of L1 protrusion for latSC photostimulation trials (cyan). Right, L1 protrusion probability (*n* = 8 mice). **c**, Photostimulation example data. Top, Experimental schematic. Photoactivation was targeted to four sites spanning 800 μm along the AP axis of the latSC (left). Middle, Still frames from an example left and right latSC photoactivation trial for each AP target at protrusion offset. Tongue tip (dot) and the tip vector (dashed line) are shown in blue, fiducial point is shown as a white dot. Bottom, Polar plots of L1 protrusion endpoints for an example left and right latSC photostimulation session. Individual colored points are individual trials, and colored dashed lines indicate the median for each AP target. Note that the L1 protrusion endpoint depended on the stimulation site. **d**, latSC photostimulation-evoked L1 protrusion angles are correlated with latSC photostimulation site (*n* = 8 mice, left). Gray lines are individual mice. Right, *R^2^* for the observed correlation between photostimulation site and L1 protrusion angle (black) compared to shuffled data (gray). **e-f**, L1 protrusion probability (**e**) and latency (**f**) did not differ between latSC stimulation sites. Data in **c-f** are color coded by stimulation site in the latSC. Colors in **c-f** correspond to latSC implant sites as shown in **c**. Colored data in **d-f** are median ± IQR. \*\*\**p* < 0.001, shuffle test; n.s., not significant, Kruskal–Wallis test. Exact statistics are in Supplementary Tables 13 and 14.

## Discussion

Humans rely on the sense of touch to guide tongue movements during eating, drinking and vocalizing^4,5^. We discovered that mice also use precise tactile feedback while drinking water from a spout. Importantly, when mice nicked a spout with the left part of their tongue surface on a straight lick, they accurately re-aimed the next lick to a left spout position. But the same nick on a right-aimed lick caused accurate re-aiming to a centered spout. Position-dependent reactions to sensory feedback require sensorimotor coordinate transformations that map postural information onto a coordinate system relevant for aiming movement. In primates, sensorimotor cortical pathways mediate touch- and visually-guided reaching and grasping^2,43,46^, and SC has been implicated in visual, auditory, and somatosensory-guided orienting^20,45,49^. Yet spinalized frogs and *Drosophila* exhibit body position-dependent responses to stimuli^44,50,51^, showing that diverse circuit architectures can implement coordinate transformations important for sensory feedback-based aiming.

Touch-guided lick aiming did not require activity in cortical areas previously associated with licking^15–17^, and instead required activity in a lateral region of the superior colliculus (latSC)^26,27^. Notably, several aspects of SC function canonically associated with other types of stimulus-guided orienting were recapitulated in our experiments. First, unilateral latSC inactivations selectively impaired contralateral touch-guided re-aiming, much like unilateral SC inactivations reduce orienting to contralateral visual targets^33,34^. Second, viral tracing identified a lingual region of the trigeminal nucleus that projected directly to the latSC, revealing an early sensory input analogous to direct retinal inputs to SC^52^. Third, latSC neurons exhibited receptive fields for mostly contralateral touch sites, and could represent these tactile events in tongue, head, and conjunctive reference frames. Fourth, latSC neural activity also encoded information about tongue position and the aiming of upcoming licks, which could arise from premotor or efference copy signals, or proprioceptive feedback. Thus, the latSC exhibits a combination of signals that are important for implementing the coordinate transformations necessary for sensorimotor feedback control^42,43,45,46^. Finally, latSC photo-microstimulation evoked licks and aimed them in accordance with a topographic map similar to that known to be used for saccades and orienting, in which more posterior stimulation sites evoke more contralaterally aimed movements^19,2019,21,38,53^. Altogether, these data show that the latSC contains a mechanosensorimotor map for touch-guided tongue control that shares mechanistic features observed in visually-guided orienting^24,25^. With recent advances examining tongue kinematics in marmosets and macaques, it will be possible to test if primate SC also implements touch-guided control^54,55^.

A comparative approach across species, behaviors, and effectors can distinguish general principles from specialized solutions to narrower control problems. The mammalian superior colliculus and analogous tectal regions in other species use sensorimotor maps to direct stimulus-guided orienting across a range of species-specific sensory modalities^19,20,38,55,58^, including whisking in rodents^25,40,41^, echolocation in bats^59^, bifoveate vision in raptors^60^, and infrared vision in snakes^61^. The topographic architecture of the SC/optic tectum is thought to implement a winner-take-all decision process important for selecting a single direction for orienting and escape^62^. Touch-guided tongue control may be similarly dependent on the SC because, like orienting, it could require a rapid selection process for both a salient sensory event and an ensuing motor response. Notably, we observed a wide range of neural signals in the latSC, including tactile responses in multiple coordinate frames, important for implementing sensorimotor transformations underlying touch-guided control. The latSC has multiple layers, cell types, and complex connectivity with brainstem, cerebellar and forebrain structures, and our findings do not rule out the possibility that other circuits may also participate in tactile-motor coordinate transformations important for tongue control. Future studies can leverage touch-guided tongue control in head-fixed mice to further clarify microcircuit-level mechanisms of coordinate transformations important for sensory-guided action^42,44,45,53^.

## Methods

### Animals and surgery

Male and female mice age 10-24 weeks old were used in this study (21 VGAT-ChR2-eYFP, JAX# 014548; 6 *Slc32a1^tm2(cre)Lowl^*^/MwarJ^ (Vgat-ires-cre), JAX # 028862; 16 C57BL/6J, JAX# 000664; 3 *Rosa^26LSL-tdTomato^* (Ai14), JAX# 007914). Mice were individually housed under a 12/12 light/dark cycle and all experiments were carried out during the dark cycle. All procedures were in accordance with NIH guidelines and approved by the Cornell Institutional Animal Care and Use Committee.

Prior to surgeries, mice were anesthetized with 5% isoflurane. Fur covering the scalp was trimmed, and mice were then head-fixed in a stereotaxic apparatus (Kopf Instruments and Leica Biosystems). An electric heat pad was used to maintain the mouse body temperature and isoflurane concentration was kept between 1-2% throughout surgeries. Mice were administered buprenorphine (0.05 mg/kg, SQ) for pain management and ocular ointment was applied to protect the eyes. The scalp was then cleaned with betadine and 70% alcohol and an incision was made along the midline to expose the skull. Enrofloxacin (5 mg/kg, SQ) and carprofen (5 mg/kg, SQ) were administered post-operatively. AP and ML stereotaxic coordinates below were taken relative to bregma, and DV was taken from the skull surface.

For optogenetic inhibition of cortical regions, craniotomies were made over ALM (2.5 mm AP ± 1.5 mm ML), TJM1 (1.76 mm AP ± 2.64 mm ML), or TJS1 (0.32 mm AP ± 3.56 mm ML)^17^ in VGAT-ChR2-EYFP mice. We implanted two 400 μm optic fibers (0.39 NA, Thorlabs) bilaterally over these areas and covered the exposed brain surface with silicone gel (Kwik-Cast, World Precision Instruments). Optic fibers were implanted with a 24 (30) degree lateral tilt in the ML/DV plane for TJM1 (TJS1). Cannulas were cemented to the skull with Metabond (Parkell). A custom-modified RIVETS headplate was then implanted over the skull^56^. We calibrated our cortical inactivations in ALM in previously published work using the same setups and laser powers used for these inactivations^11^.

For optogenetic inhibition of SC through the SNr-SC pathway, 400 nL of AAV5-EF1a-double floxed-hChR2(H134R)-EYFP (Addgene) was bilaterally injected into SNr (−3.2 mm AP ± 1.5 mm ML, −4.3 mm DV) of VGAT-IRES-Cre mice, and two 100 μm optic fibers (0.22 NA, Doric Lenses) were implanted bilaterally over the intermediate layers of the SC (−3.4 mm AP, ± 1.5 mm ML, −2.0 mm DV) along with a headplate. We waited at least 4 weeks for viral expression before performing any manipulations, during which time mice underwent behavioral training.

For photostimulation of SC, 500 nL of AAV2/hSyn-hChR2(H134R)-EYFP-WPRE-PA or AAV5/hSyn-hChR2(H134R) (UNC Vector Core, a gift from Karl Deisseroth) was unilaterally injected into SC (−3.4 mm AP, ± 1.5 mm ML, −2.6 mm DV) of C57BL6/J mice. Four fiducials were made with marker ink around the craniotomy for later calibration. The craniotomy was then sealed with silicone gel, and a headplate was implanted with a thin layer of clear Metabond covering the skull surface.

For acute electrophysiology experiments, fiducials were made bilaterally over ALM and/or SC, as well as visual cortex (−2.3 mm AP, ± 3.0 mm ML), and marked with black ink. Additional fiducials over bregma and lambda were also made for later calibration. A headplate was then implanted with a thin layer of clear Metabond covering the skull surface.

For cortical lesions, fiducials were marked bilaterally over ALM, TJM1, and TJS1, with additional four fiducials for calibrating skull leveling later. A headplate was then implanted with a thin layer of clear Metabond covering the skull surface. After mice were trained in the behavioral task and pre-lesion data was collected, the clear Metabond covering the cortical areas was removed and re-sealed with silicone gel. About 24 hours later, 125 nl of 4% N-Methyl-D-aspartic acid (Sigma Aldrich M3262, diluted in 1X DPBS (Corning) with 1M NaOH) was injected into each site in ALM (1.0 mm and 0.6 mm below the brain surface), TJM1 (1.0 mm and 0.6 mm below the brain surface), and TJS1 (0.32 mm AP ± 3.45 mm ML, 0.6 mm below the brain surface; 0.32 mm AP ± 3.10 mm ML, 1.0 mm below the brain surface). We decided to perform the surgeries on two separate days to ensure the survival of animals by reducing the length of anesthesia on a single day. Additionally, supplementary water and Nutra-Gel (Bio-serv) was given to animals from 1 day before the surgeries to 1-3 days after depending on recovery status indicated by weight.

### Viral tracing

2-3 μL of AAV1-hSyn-Cre was injected into the tongues of Ai14 reporter mice to label processes of primary trigeminal afferents that innervate the tongue. Additionally, 100 nL of AAV5-CAG-GFP was injected dorsally into SpVO to label secondary trigeminal neurons. Mice were sacrificed after ∼8-12 weeks. AAV1-hSyn-Cre was a gift from James M. Wilson (Addgene viral prep #105553-AAV1; http://n2t.net/addgene:105553; RRID: Addgene_105553). AAV5-CAG-GFP was a gift from Edward Boyden (Addgene viral prep #37825-AAV5; http://n2t.net/addgene:37825; RRID: Addgene_37825).

### Histology

Animals were sacrificed with a lethal dose of IP sodium pentobarbital, followed by pericardial perfusion with 4% paraformaldehyde (Electron Microscopy Services) in 1X PBS (Corning). Brains were extracted, sliced to 50 or 100 μm thickness with a Leica VT1000 S vibratome, mounted on glass slices with PVA-DABCO (Sigma Aldrich), and then imaged with a Zeiss LSM 710 confocal. To validate cortical lesions, brain slices were first incubated in 0.6% Triton in 1X PBS for 20 minutes, washed in 1X PBS (5 minutes × 3 times), stained with mouse anti-NeuN antibody conjugated with Alexa Fluor 555 (EMD Millipore MAB377A5, 1:250 dilution in 1X PBS with 0.3% Triton) for 1.5 hours, and finally washed again in 1X PBS (5 minutes × 3 times).

### Behavior

The behavioral setup was as previously described^11^. Briefly, the tongue was captured from side and bottom views by placing a mirror (Thorlabs ME1S-P01 1′′) at 45° underneath the face of the mouse. Videos were acquired at 1,000 fps with a Phantom VEO 410L camera and a Nikon 105-mm f/2.8D AF Micro-Nikkor lens. An 880 nm wavelength IR strobe (AOS Technologies, manufacturing discontinued) or 850 nm wavelength IR ring light (Smart Vision Lights) was used to provide illumination of the tongue during video imaging. To build the behavioral rig, we 3D-printed custom-made head clamps for fixation. 1 kHz audio cues were played through tone generators (Med Associates), and a blue LED (Amazon) optogenetic masking light was delivered throughout the entire trial period. Water was dispensed through a 0.072 in outer diameter stainless steel spout (Small Parts) using a 24- or 5-volt solenoid valve (The Lee Company). Spout contacts were detected either with a capacitive touch sensor (Atmel), or a custom-made electrical lick sensor^57^ to increase contact detection sensitivity and reduce contact-induced noise in electrophysiology recordings. For double-step experiments, water spouts were attached to two orthogonally-arranged linear motors (Faulhaber) mounted on guide rails (McMaster-Carr). Control signals sent to the controllers for the servo motors (Faulhaber) were recorded throughout the experiment, allowing us to track the spout during behavioral sessions (see section Tongue contact location tracking). During experiments, water spouts were displaced by a pre-calibrated distance along both AP and ML axes. Custom LabVIEW code for behavioral tasks were run using NI sbRIO-9636 or −9637 FPGAs.

### Behavioral paradigms and training

Water deprivation started five days post-surgery. Mice under water deprivation received at least 1 mL of water daily from performing behavioral tasks. Supplementary water was provided to meet this requirement if mice did not collect enough water from tasks, or on days when mice did not take part in any behavioral testing. After the body weight of mice stabilized (typically around 80% of the original weight), mice began behavioral training following a similar procedure as previously described^11^. Mice were first habituated to head-fixation on the behavioral rig for three days. Then, mice began cued-lick training in a trial structure. To encourage mice to lick the water spout, water was dispensed at cue onset on the first few sessions. Once mice reliably began to associate the auditory cue with a water reward (typically, after one to three sessions), we made the water dispense contingent on the first spout contact within 1.3 s of the auditory cue. We then imposed a 1 s no-spout-contact window before the audio cue to discourage premature licking prior to cue onset. Any spout contact during this window ended the trial and initiated a new inter-trial interval. The inter-trial interval was randomly chosen from an exponential distribution with a flat hazard rate. Once mice could reliably trigger water dispense for over 95% of trials in a session with very few licks during the inter-trial interval and less than 10% no-spout-contact window violations, we either started the photo-microstimulation experiment or moved on to the next phase of training for the left/right doublestep paradigm.

For all analyses, with the exception of photostimulation experiments, lick numbers (L1, L2, etc.) are defined relative to the first lick that made contact with the spout after the auditory cue on a given trial. For photostimulation experiments, lick numbers were defined relative to stimulation onset. For assessing the kinematic impairments in cortical lesioned mice (Extended Data Fig. 4), the first lick was defined as the first lick after auditory cue onset regardless of spout contact.

For left/right displacement experiments, the water spout was subtly (∼1.1 mm) displaced to the left or right at L1 contact offset (left, center, right positions each with 33% probability). For left or right positions the spout was also moved slightly closer in the AP direction such that the radial distance from the jaw to the spout was similar across all possible spout positions. Except for the recentering experiment described below, the spout remained at the same position for the remainder of the trial following the displacement and only moved back to the center position one second prior to the start of the next trial. To ensure that mice could not detect spout movements by seeing, whisking or hearing spout movements, we placed the water spout ∼3 mm beneath the snout out of the visual field and trimmed the whiskers and orofacial hair of mice every 2-4 days in all experiments. The auditory cue, which plays throughout the duration of the trial, served as a masking sound for any residual noise made by the servo motors, which were already quiet. Water was dispensed on L2 contact or L3 contact (Extended Data Fig. 1). Mice were trained with this new task setup for 7-14 days, with most mice already exhibiting at least some L3 re-aiming on day 1 (data not shown). For the first few days of training, we only allowed two consecutive spout displacements to the same direction to ensure mice did not develop a directional licking bias on L2. This limit was increased to ten during experiments.

For recentering experiments, water spouts were randomly (∼35% probability) moved back to the center position at L3 contact offset. For 3 out of 5 mice, the recentered position was slightly (∼0.5 mm) closer than the initial center position to ensure mice would still nick the spout with lateral tongue protrusions on L4. Recentering trials first began 1-3 days prior to experiments to familiarize mice with the task. During training, we lowered the probability of recentering to only 15-20% to prevent mice from potentially developing a center protrusion bias on L4.

For cortical lesion experiments, two out of five animals exhibited such pronounced hypometria that they were unable to reach the 3 mm spout distance used in other experiments. We anticipated this and collected pre-lesion data at various spout distances, including approximately 2.25 mm, which all animals were able to reach 6-16 days post-lesion. Data from all mice in Extended Data Fig. 4 were collected at this 2.25 mm distance.

### Optogenetic inactivation

We used laser diode light sources (LDFLS_450-450, Doric Lenses), attached to an optical rotary joint splitter (FRJ_1×2i_FC-2FC_0.22, Doric Lenses) and delivered light bilaterally or unilaterally to the implanted optic fibers using 200 μm, 0.22 NA or 400 μm, 0.43 NA, lightly armored metal-jacket patch cords (Doric Lenses). The light sources were set to analogue input mode and driven with a square pulse for 250 ms with a 50 ms ramp down starting at L2 contact onset, which was detected in real time. For cortical photoinactivations, laser power was set to 10 mW. For bilateral SC photoinactivation, laser power was set to 5 mW. For unilateral SC photoinactivation, most animals were inactivated with 5 mW. However, one animal was inactivated with 1 mW power. We observed no difference in the magnitude of the effect of unilateral manipulations between the 1 and 5 mW cohorts (data not shown), and thus combined the data. Pulse duration and laser powers were identical for photovalidation experiments.

### Optogenetic stimulation

Light was delivered at 1 mW with laser diode light sources through a 200 μm, 0.22 NA lightly armored metal-jacket patch cord (Doric Lenses) to a very thin silica optic fiber (50 μm diameter, 0.22 NA, Doric Lenses). Simulations performed by Doric Lenses with a scattering model for the same optic fiber parameters predicted that this would illuminate a volume underneath the fiber constrained approximately within 50-70 μm in the AP and ML direction and extending at least 250 μm in the DV direction (incident flux around 100 mW/mm^2^ at 250 μm underneath the tip). 12-24 hours prior to the start of experiments, the thin layer of Metabond covering the craniotomy was removed and re-sealed with a layer of silicone gel. During the experiment, the optic fiber was held in place with a custom-made fiber cannula holder and targeted stereotaxically to different sites of the latSC (−3.1 to −3.9 mm AP, ± 1.3-1.6 mm ML for −3.1 mm AP and ± 1.6-1.8 mm ML for all other sites). The DV coordinate for each site was adjusted between −1.75 to −2.45 mm for each site such that the fiber tip sat above the intermediate layer of SC. AP and ML coordinates were calibrated based on previously marked fiducials centered at bregma and lambda, and DV coordinates were based on micromanipulator readings relative to the bregma skull surface. Following acute fiber implantation, the craniotomy was filled with silicone gel to reduce brain movement and 1X PBS was pipetted into the headplate to prevent the brain from drying. For each site, laser light was first delivered at cue onset for 250-400 ms (adjusted to ensure laser was on for at least the entire duration of L1) with ∼45% probability for ∼120 trials, then for ∼30 trials without auditory cue to indicate trial structures. One to three sites were tested with this sequence on each day, after which the craniotomy was re-filled with silicone gel. On the last day of the experiment, thin needles (NF36BV WPI) were coated with a lipophilic dye (DiI or DiD, Thermo Fisher) to mark all previously stimulated sites.

### Acute electrophysiology

Extracellular recordings were made acutely using 64-channel silicon probes (ASSY-77 H6, Cambridge Neurotech). For latSC photovalidation experiments, a 200 μm optic fiber was attached to the silicon probe (ASSY-77 H6 with 200 μm fiber, Cambridge Neurotech). The 64-channel voltage signals were amplified, filtered and digitized (16 bit) on a headstage (Intan Technologies), recorded on a 512-channel Intan RHD2000 recording controller (sampled at 20 kHz), and stored for offline analysis. At 12–24 h before recording, a small (1.5 mm diameter) craniotomy was made unilaterally over SC, as well as visual cortex. A ground screw with a soldered gold pin (A-M systems) was implanted into the skull overlying visual cortex away from the recording site and held in place with Metabond. The probes were targeted stereotaxically to SC, lowered to a depth of 2.1-2.6 mm below the dura surface. Recording sites in the AP plane spanned 1 mm from −2.9 to −3.9 mm AP. The ML recording location was targeted to the lateral region of the SC at each AP site, which changed with the contour of the SC along the AP axis from −1.2 mm ML at more anterior sites to −1.6 mm ML at more posterior sites. For cortical photovalidation experiments, probes were targeted to the deep (−0.8 to −1.2 mm DV) or superficial (−0.3 to −0.8 mm DV) layers of ALM (−2.5 mm AP, ± 1.5 mm ML) and a 400 μm optic fiber was placed on the dura surface. Photovalidation recordings were only performed in ALM as photoinactivation methods were identical for all three cortical regions investigated (ALM, TJM1, and TJS1). AP and ML coordinates were calibrated based on the fiducials previously marked and DV coordinates were based on micromanipulator readings. Following probe insertion, 2% low-melt agarose (A9793-50G, Sigma Aldrich) in 1X PBS was pipetted in the craniotomy to reduce brain movement and additional PBS was pipetted into the headplate to prevent the brain from drying. Three to seven recordings were made from each craniotomy. At the end of each session, the craniotomy was covered with silicone gel. Probes were coated with a lipophilic dye (DiI or DiD, Thermo Fisher) prior to each recording to ensure placement in the region of interest.

### Lingual Kinematics Analysis

#### Lingual segmentation

We implemented a semantic segmentation neural network (U-NET) to identify and segment the tongues from high-speed videography as previously described^11^. Our network was trained on 7,681 frames randomly selected from 36 mice. Training data for tongue segmentation were hand-traced using a custom MATLAB GUI. We trained networks for side and bottom views respectively in batches of 256 images, using the ‘adam’ optimizer and a binary cross entropy loss function. The networks were trained until the loss function reached an asymptotic value of 0.0035 for the side-view network and 0.0036 for the bottom-view network, with a validation accuracy of 0.9984 and 0.9985, respectively. Both networks reached asymptotic performance within 5,000 epochs.

#### Lingual kinematics analysis

The full 3D kinematics of the tongue tip from high-speed videography were obtained as previously described^11^. Briefly, we performed a visual hull reconstruction from the side and bottom view silhouettes of the tongue. The centroid of the tongue was identified from the 3D voxels and a two-step search process was implemented to locate the tongue tip. Tongue tip kinematics were smoothed using an 8-pole, 50 Hz low-pass filter. Lick phases were defined based on rate of volume changes calculated from the 3D tongue reconstruction as the following: 1) protrusion phase as from first moment of tongue detection to the first volume change rate minimum; 2) retraction phase as from the last volume change rate minimum to the last moment of tongue detection; 3) CSM phase as from protrusion phase offset to spout contact onset; 4) SSM (spout submovement) phase as from spout contact onset to retraction phase onset.

For kinematics variables, we calculated speed as the magnitude of the one-sample difference of the position vector, acceleration as one-sample difference of speed, and pathlength by integrating the speed over the entire lick duration,. Number of acceleration peaks were determined using MATLAB’s findpeaks function. Protrusion angle was defined as the angle between the facial midline and a vector pointing from a fiducial point on the jaw to the tongue tip (Fig. 1c). The facial midline was defined as the line that passes through the midpoint between the nostrils and the midpoint between the incisors. The fiducial lies on the midline and was calibrated for each individual mouse to be at approximately the same distance posterior to the mouth, aligned in the AP direction with the visible base of the jaw.

When examining the kinematics of any lick, we excluded trials where the mouse contacted the spout twice or missed the spout on any lick prior to the lick being analyzed. To calculate the correlation between L2 contact location on the tongue and the protrusion angle change on the next lick, we first calculated the protrusion angle change (the difference in protrusion angle from L2 to L3 for all trials, Extended Data Fig. 7a). We then correlated the L2 contact angle with the L2 to L3 protrusion angle change across all conditions (Extended Data Fig. 7c), as well as for each condition separately (Extended Data Fig. 7d-f).

#### Tongue contact location tracking

To create a 3D model of the spout, we first identified all of the discrete spout positions (e.g., left, center, right) used in a single behavioral session using example frames from videos of that session. We then used recorded command voltages sent to the servo motors to infer the 3D location of the spout tip in all other video frames from that session. Given the 3D spout location and the known size (0.072 inch outer diameter) and shape (cylinder) of the spout, we generated a full voxel representation of the spout for each frame.

For these analyses, we used spout contact sensor data to identify the set of video frames corresponding to the entire duration of the tongue-spout contact event on each lick. To identify which parts of the tongue surface contacted the spout, we first embedded the 3D representation of the spout in the same space as the 3D visual hull reconstruction of the tongue. We then performed an iterative search to identify which tongue voxels (each 60 μm^3^) were either overlapping with, or one voxel away from, spout voxels; these were defined as contact voxels. We then took the centroid of these contact voxels and defined this as the contact centroid. This was repeated for each timestep throughout the entire duration of tongue-spout contact for all licks that made contact with the spout. Next, we identified a similar facial landmark on the face of each mouse as a fiducial point. To estimate contact site on the tongue surface, we were mindful that the tongue is a highly deformable object without useful landmarks, and therefore Euclidean positions on a fixed surface do not exist in the same way as for a finger or body, for example. Thus we chose to quantify the contact site as a single variable, the contact angle. To obtain the contact angle for each lick, we calculated the angle between the tongue-tip vector (vector pointing from the fiducial to the tongue tip) and the contact centroid vector (vector pointing from the fiducial to contact centroid). We chose a polar coordinate system because it captures in a single angular measurement the variance in contact site in both mediolateral and anteroposterior axes that were elicited in our task. Note also that this method can estimate the contact site at every timestep during spout contact (i.e., a contact trajectory); however, for analysis we take the contact angle at contact onset to reduce the trajectory to a single point. To assess head-centric tuning, we calculated head-centric contact angle as the angle between the contact centroid-vector and the midline of the face (defined in ‘Lingual kinematics analysis’ section above).

#### Analysis of optogenetic manipulations on lick kinematics

For bilateral and unilateral optogenetic manipulations, we analyzed differences in L3 protrusion angle (Fig. 1f & 2e), and all other kinematic parameters (Extended Data Fig. 2e-j & Extended Data 5a-f) between no laser and laser conditions using a shuffle test. For unilateral optogenetic manipulations, we additionally calculated the angular correction magnitude by taking the difference between the center spout trial L3 protrusion angle and the contralateral or ipsilateral spout trial L3 protrusion angles for laser on and laser off trials. The difference between laser on and laser off trials was significant if the *p*-value was less than 0.025 on either tail. Additionally, we compared the effect inactivating each brain region (ALM, TJM1, or latSC) had on correction magnitude for ipsilateral or contralateral corrections separately, using a shuffle test. We considered these as two sets of multiple comparisons, examining the *p*-value against the Bonferroni corrected *p*-value (*p* < 0.05/(3×2) = 0.008, three pairs of comparisons, tests on both tails).

For optogenetic stimulation experiments, we defined the protrusion angle for the first lick evoked by stimulation as the angle between the facial midline and a vector pointing from the fiducial point to the tongue tip at the moment of maximum tongue tip lateral displacement. We excluded data from sites that histology indicated may be outside the latSC or outside regions with viral expression (Extended Data Fig. 10a). We combined data with the same AP coordinates for each mouse. We then ran a correlation test between the AP coordinate of microstimulation sites and the median L1 protrusion angles for each animal, the results of which we report in Pearson’s *R^2^*. We then asked whether the observed *R^2^* value was due to chance by performing a shuffle test, where the null distribution was created by shuffling condition (AP site) with L1 protrusion angle. These analyses were repeated for cue-only, stimulation-only and stimulation-with-cue experiments. To analyze L1 protrusion probability and latency across stimulation sites, we ran the Kruskal–Wallis test with Tukey’s HSD post-hoc test. We also analyzed the differences between kinematic variables across cue-only, stimulation-only, and stimulation-with-cue conditions using a shuffle test examining the *p*-value against the Bonferroni corrected *p*-value (*p* < 0.05/(3×2) = 0.008, three pairs of comparisons, tests on both tails). For analysis of L1-L2 inter-lick interval, we only included trials where L1 did not make spout contact and L2 was initiated during stimulation.

### Electrophysiology analysis

#### Spike sorting and unit selection criteria

Extracellular voltage traces were first notch-filtered at 60 Hz. The data were then spike-sorted automatically with Kilosort2 (https://github.com/MouseLand/Kilosort)^58^, and curated manually with Phy2 (https://github.com/cortex-lab/phy). During manual curation, units containing low-amplitude spikes and/or non-physiological or inconsistent waveform shape were discarded and not included in further analyses. Neurons with fewer than 10 trials in any of the conditions tested were excluded for all analyses performed below. Additionally, neurons that fired less than 0.1 Hz, either throughout the entire trial period or within a given time window defined in the sections below, were excluded from further analysis.

#### Rate histograms

The rate histograms for example neurons were computed by binning the spiking activity of each example neuron in 10 ms bins and aligning activity to trial onset (Extended Data Fig. 3), L2 contact (Fig. 3a, Fig. 4c & d, Extended Data Fig. 6a & 8), L4 contact (Fig. 4c & d, Extended Data Fig. 8), or L4 protrusion onset (Extended Data Fig. 9a & e). For principal component analysis of neural data, 5 ms bins were used to allow for a better estimate of divergence time and activity was aligned to L2 contact onset. Activity was smoothed by taking the moving average of the firing rate for each neuron in the preceding 3 time bins. To create rate heatmaps across selectively modulated neurons, rate histograms for all selectively modulated neurons were computed as described above, and data were aligned to L2 contact onset (Fig. 3b, Extended Data Fig. 6b). Selectivity for each neuron was determined as described below in the section ‘Single-unit significance testing.’ For each selectivity heatmap, only neurons that were selectively modulated by a specific contact site (contralateral, center, etc.) were chosen for display. For all conditions within each selectivity heatmap, neurons were sorted from top to bottom by the time of their peak mean firing rate within a 100 ms analysis window following L2 contact in the condition it was selective for. The firing rate of each neuron was min-max normalized in a window ± 300 ms around L2 contact. Note that, for some neurons, this means that the absolute peak firing rate for the neuron in the ± 300ms window may have occurred outside of the selectivity analysis window. For photovalidation heatmaps, rate histograms for each condition (laser on/off) were instead aligned to laser onset (Extended Data Fig. 3), min-max normalized, and sorted in ascending order from top to bottom by the mean firing rate within the laser on condition. Only neurons with a mean firing rate of > 1 Hz within the analysis window on non-laser trials were included.

#### Single-unit significance testing

To determine whether the firing rate of a neuron was significantly modulated by the trial condition (left, center, right) following L2 contact (Fig. 3a-c, Extended Data Fig. 6a-b), we calculated the firing rate in a 100 ms time window following L2 contact onset for each trial. We then performed a one-way analysis of variance (ANOVA) to test whether the firing rate of each neuron was modulated by condition. If a given neuron was significantly modulated by condition using ANOVA (*p* < 0.05), we then additionally performed pairwise post-hoc tests (Tukey’s HSD) to determine what condition pairs were significantly modulated with each other (left/right, left/center, center/right). A condition pair was significantly modulated if the *p*-value was less than the Bonferroni corrected *p*-value (*p* < 0.05/3). A neuron was said to be selective for a given condition if the mean firing rate for a given condition was significantly greater than the mean firing rate in all other conditions. A neuron was said to be selective for two conditions if the mean firing rate for two conditions were significantly greater than the third, but those two conditions were not significantly different from one another.

To test whether the relative proportion of selective neurons differed by AP recording site in the latSC (Extended Data Fig. 6c), we recorded the estimated AP/ML recording location within latSC during each recording session in a subset of our animals (*n* = 4 animals) using micromanipulator readings. Silicon probes were coated with either DiI or DiD on each session to verify the recording site. The specific dye used was alternated on each session. We then grouped neurons based on their recording location into 5 bins, spanning from −3.0 to −3.8 (± 0.1) mm AP in latSC, in 0.2 mm increments (e.g., −3.0 mm AP includes sites spanning −2.9 to −3.1 mm AP). To test whether the number of neurons selective for one particular condition was greater in proportion relative to other conditions, we performed a bootstrap test. We randomly sampled with replacement neurons from each recording site and assessed the proportion of selective neurons for each condition on each iteration for 100,000 iterations. The proportion of neurons selective for a particular condition (contra, for example) was said to be significantly greater than another condition (ipsi, for example) if the number of iterations where the proportion of neurons selective for the other condition (ipsi) was greater than the condition of interest (contra) was less than the Bonferroni-corrected *p*-value (*p* < 0.05/10 = 0.005).

To test whether a neuron was modulated during recentering trials on L4 (Extended Data Fig. 8b-f), we calculated the firing rate in a 100 ms time window following L4 contact onset for each trial. As recentering trials only occurred on left/right displacement trials, we performed two *t*-tests; one between left and left recentering trials, and another between right and right recentering trials. A neuron was modulated on L4 if either one of these tests had a *p*-value less than the Bonferroni corrected *p*-value (*p* < 0.05/2).

#### Contact location information of latSC neuronal activity

For these analyses (Fig. 3d), we excluded neurons with a mean firing rate of less than 3 Hz within the analysis window (*n* = 422/553 neurons passed this criteria). To determine the amount of contact location information (in bits/spike) present in individual latSC cells, we repurposed analyses previously used to analyze spatial information in hippocampal place cells^39^. Briefly, contact location information was obtained from the following equation: 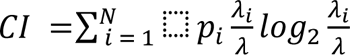, where *N* is the total number of contact angle bins, *p_i_* is the probability of occupancy, *λ_i_* is the firing rate in the *i*th contact angle bin, and *λ* is the mean firing rate of the neuron. To test whether a neuron was significant for contact location information, we performed a shuffle test. For each neuron, we randomly permuted contact angle relative to their respective binned firing rates 1000 times and asked whether the observed contact location information value was outside of the 95th percentile of the shuffled distribution.

#### Correlating latSC neural activity with contact angle

We only included neurons with significant L2 selectivity for L2-only correlation analyses (Fig. 3a & e, Extended Data Fig. 6a), only neurons with significant L2 and L4 selectivity for recentering analyses (Fig. 4i, Extended Data Fig. 8c). We also excluded neurons with a mean firing rate of less than 3 Hz in the analysis window for L2 correlation analyses (*n* = 422/553 neurons passed this criteria). For neurons that passed this criteria, L2 (or L4) contact angles were binned by 3° (contact bins) and the mean firing rate following L2 contact in each contact bin was calculated. We required at least 3 trials in each bin for that bin to be included in the analysis. A correlation was considered significant if the actual R^2^ is greater than the 95th percentile of *R^2^* values obtained by permuting the firing rate with respect to contact angle over 1,000 iterations. We calculated head-centric contact angles as the angle between the contact vector and the facial midline (which passes through the fiducial), both in the bottom view.

#### Principal component analysis (PCA) of latSC population activity

To obtain an estimate of when neural activity on left and right trials diverged from center trials (Fig. 3f), we first used PCA to reduce the dimensionality of our data as previously described^11,59^. Data were aligned to L2 contact onset, and peri-stimulus time histograms (PSTHs) were generated in 5 ms bins to create a matrix of (neurons x time bins x conditions). We then ran PCA on this data and projected the condition-averaged (left, center, and right) responses onto the first 19 dimensions of this space, which together explained >90% of the neural variance in our dataset. The neural distance was calculated as the Euclidean distance between the left and center or right and center trajectories in the first 19 dimensions. We plotted the trajectories from each condition in the first three dimensions of this space (Fig. 3f, left panel), in addition to the median times of behavioral variables, such as the time of L2 contact, on these trajectories.

To obtain the time of divergence between trajectories, as well as an estimate of the variability of these distances, we performed a hierarchical bootstrap. For each condition (left, center, and right), we first resampled with replacement neurons, and then resampled with replacement trials for each condition. We then computed PSTHs with this resampled dataset, projected the data onto the top 12 principal components, and again calculated the Euclidean distance between the left and center and right and center trajectories. This procedure was repeated 1,000 times to yield the bootstrapped estimate of the standard deviation of the distance between these trajectories. The time of divergence between neural trajectories was then defined as the median time that the neural distance exceeded two standard deviations above the mean baseline activity (−300 to 0 ms before L2 contact onset) across both left/center and right/center bootstraps.

#### Contact response latencies of single latSC neurons

For determining the L2 response latencies of latSC neurons, we restricted our analyses to only neurons with significant L2 selectivity. For each neuron, we aligned neural activity to L2 contact onset in 1 ms bins and performed Wilcoxon rank-sum (WRS) tests on the number of spikes in 5 ms windows across: 1) left vs. right, 2) left vs. center, and 3) center vs. right conditions, similar to previous analyses used for estimating neuronal response latencies to distorted auditory feedback in singing birds^60^. Windows were shifted in 3 ms increments from 0 - 250 ms relative to L2 contact onset. The response latency was defined as the first bin where at least 3 consecutive bins had a Bonferroni-adjusted *p*-value of < 0.0055 = (0.05/(3×3), 3 conditions, 3 bins).

To examine the response latency of latSC neurons depending on coordinate frame (tongue-centric, etc.), we first determined the coordinate frame (or lack thereof) of each latSC neuron by correlating neural activity following L4 contact with contact location in either a tongue- or head-centered reference frame, as described in section ‘Correlating latSC neural activity with contact angle’ above. For these analyses, we only included neurons recorded during recentering sessions and with significant L2 and L4 selectivity. We then calculated the median response latency across all neurons of a given reference frame and performed a hierarchical bootstrap over 1,000 iterations for significance testing and to get an estimate of the interquartile range. Response latencies in a head-centric reference frame were considered to be significantly longer than latencies in another reference frame if the *p*-value was lower than the Bonferroni-adjusted *p*-value of < 0.016 (0.05/3).

#### Lick cycle cross-correlation analysis

To determine whether the activity of a neuron was correlated with lick cycle (Extended Data Fig. 9a-c), we analyzed the time window from 50 ms before L4 protrusion onset to 300 ms after L4 protrusion onset in non-recentering trials only. We chose L4 for this analysis because the spiking of latSC neurons prior to L4 was often dominated by L2 contact and L3 re-aiming responses, and we chose non-recentering trials to ensure recentering contacts would not similarly dominate L4 responses. We calculated the average firing rate and tongue volume across trials with a 10 ms bin size for left, center and right trials separately. Only bins with at least 10 trials were included in this analysis. Neurons with a firing rate lower than 5 Hz during the analysis window were excluded. Cross-correlation was then performed between the mean-subtracted firing rate of a neuron and tongue volume with normalization. The lag was determined as when the maximum cross-correlation coefficient was obtained within a ± 100 ms lag window (Extended Data Fig. 9d). For each trial type, a neuron was considered significantly correlated with the lick cycle if the 1st percentile of the bootstrapped maximum cross-correlation coefficient values was greater than the median shuffled maximum cross-correlation coefficient values obtained by randomizing spike time within each trial. Both the bootstrap and randomization were performed over 1,000 iterations.

#### Encoding model analysis

We used Poisson GLMs to determine whether the activity of single latSC neurons encoded tongue position information prior to L4 contact (Extended Data Fig 9f-i). Note that, we are using GLMs as an analysis method to examine coding of tongue position in the latSC, not as a model of the latSC, per se. Inspired by previous work^47,48^, we sought to make predictions of latSC neuronal activity on single licks, rather than across an arbitrary time window prior to L4 contact. We summed the number of spikes on each trial from 50 ms prior to L4 protrusion onset until L4 contact onset and computed two families of features for each protrusion: tongue tip position at L4 protrusion offset (x/y/z components) and previous (L3) spout contact location (x/y/z components). Although we were most interested in testing whether tongue position was encoded in pre-contact L4 latSC neuronal activity, given the close temporal proximity (∼120 ms) between past (L3) contact and L4 re-aiming, any relationship we might observe between L4 tongue position and latSC neuronal activity could be confounded by past L3 contact location. We included both variables to examine whether L4 tongue position was better encoded than past L3 contact location in latSC pre-contact L4 activity. We also included two other predictors: protrusion duration prior to L4 contact and an estimate of firing rate drift over the session. Protrusion duration prior to L4 contact was included, since any difference in duration between trial types could give rise to spurious correlations with tip position. We estimated drift by dividing the trials over a session into 5 equally sized chunks and computing the mean firing rate in each chunk. We took the log of both predictors to account for the exponential nonlinearity. For these analyses, only neurons with a firing rate greater than 5 Hz during the analysis window were included (*n* = 320/553 neurons).

We fitted GLMs with the ‘fitglm’ function in MATLAB, using an exponential link function. We used five-fold cross-validation to assess the goodness-of-fit of our models. Goodness-of-fit was assessed using McFadden’s pseudo-R^2^, which was calculated on held-out test data on each fold of cross-validation and averaged across all folds. Note that pseudo-R^2^ values of 0.2 - 0.4 represent excellent fits.

We included in our analyses a null model that consisted only of protrusion duration prior to L4 contact and firing rate drift as predictors. To test for significance between the full and null model, we first calculate the likelihood ratio test statistic in the observed data. We then calculated this same statistic on 1,000 instances of shuffled data, where the rows of the design matrix were permuted relative to spiking activity. A neuron was included in further analysis if the probability of the shuffled likelihood ratio being greater than the actual likelihood ratio was less than a *p*-value of 0.05. To test for significance of each variable, we repeated the above analysis, except between the log-likelihood of the full model and each partial model. A neuron was considered significant for a predictor if the probability of the shuffled likelihood ratio being greater than the actual likelihood ratio was less than the Bonferroni-corrected *p*-value (*p* < 0.05/6).

The relative contribution of each predictor for each neuron was calculated as previously described^47,48^ from the following equation: 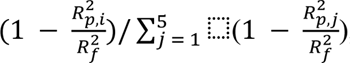, where **R*^2^_p,i_* is the variance explained by the partial model excluding the *i*th predictor and **R*^2^_f_* is the variance explained by the full model. Relative contributions were averaged across all neurons to create Extended Data Fig. 9h.

#### Statistics

*t*-tests, correlation, linear regression, Kruskal-Wallis, likelihood ratio, and ANOVA tests were performed with built-in functions of MATLAB. Shuffle tests were used for behavior, and photomanipulation analyses. We performed hierarchical bootstrapping where required by re-sampling mice with replacement, then resampling trials with replacement within each condition. The parameter (mean, median, etc.) or statistic (correlation coefficient, etc.) of interest was then calculated on each round of bootstrapping.

## Supporting information

Supplementary Tables 1- 14. This file contains Supplementary Tables 1-14, which include statistics for each figure panel.

Mice use tactile feedback to re-aim licks. Side and bottom views of the mouse tongue for L1-L5 at full speed and slowed 1/40x, for left, center and ri

Bilateral ALM, TJM1, or TJS1 inactivation does not impair touch-guided re-aiming. Example trial in the touch-guided lick task with bilateral ALM, TJM1

Lesions of all three lingual cortical areas at once impair lick kinematics while preserving touch-guided re-aiming. Example pre-lesion and post-lesion

Bilateral latSC inactivation halts ongoing licking. Example trial in the touch-guided lick task with bilateral latSC photoinactivation from L2 contact

Unilateral ALM or TJM1 inactivation biases licks ipsilaterally, but preserves touch-guided re-aiming. Example trial in the touch-guided lick task with

Unilateral latSC inactivation impairs contralateral touch-guided re-aiming. Example trial in the touch-guided lick task with unilateral latSC photoina

Recentering task decouples contact site from spout position. Example trial from a recentering session. In addition to left or right displacements betw

latSC photostimulation reveals a topographic map for lick aiming. Example trials with unilateral latSC photostimulation at 4 sites along the AP axis i

## Data Availability Statement

Data will be made available upon request.

## Code Availability Statement

Custom MATLAB and Python code used for analysis is available on Github here: https://github.com/GoldbergLab

## Acknowledgements

We thank Chris Rodgers for advice on generalized linear models. We thank Melissa Warden for Zeiss LSM 710 confocal use. We thank Teja Bollu, Joe Fetcho, Andrew Pruszynski, Michael Sheehan, Melissa Warden and Goldberg lab members for helpful discussions. We thank Teja Bollu, Antonio Fernadez-Ruiz, Joe Fetcho, Gaby Maimon, Azahara Oliva, Madineh Sedigh-Sarvestani and Matthew Zipple for comments on the manuscript.

## Author Contributions

These authors contributed equally: Brendan S. Ito, Yongjie Gao. B.S.I, Y.G. and J.H.G. designed the experiments. B.S.I and Y.G. conducted all experiments, analyzed all data and trained all animals. B.S.I. and B.M.K. developed the contact location tracking algorithms. B.S.I., B.M.K. and Y.G. built both the hardware and software necessary for data acquisition. B.S.I. and J.H.G. wrote the manuscript.

## Competing Interest Declaration

The authors declare no competing interests.

## Supplemental Information

Supplementary Tables 1-14.

Supplementary Videos 1-8.

## Extended Data Figures

**Extended Data Fig. 1.**
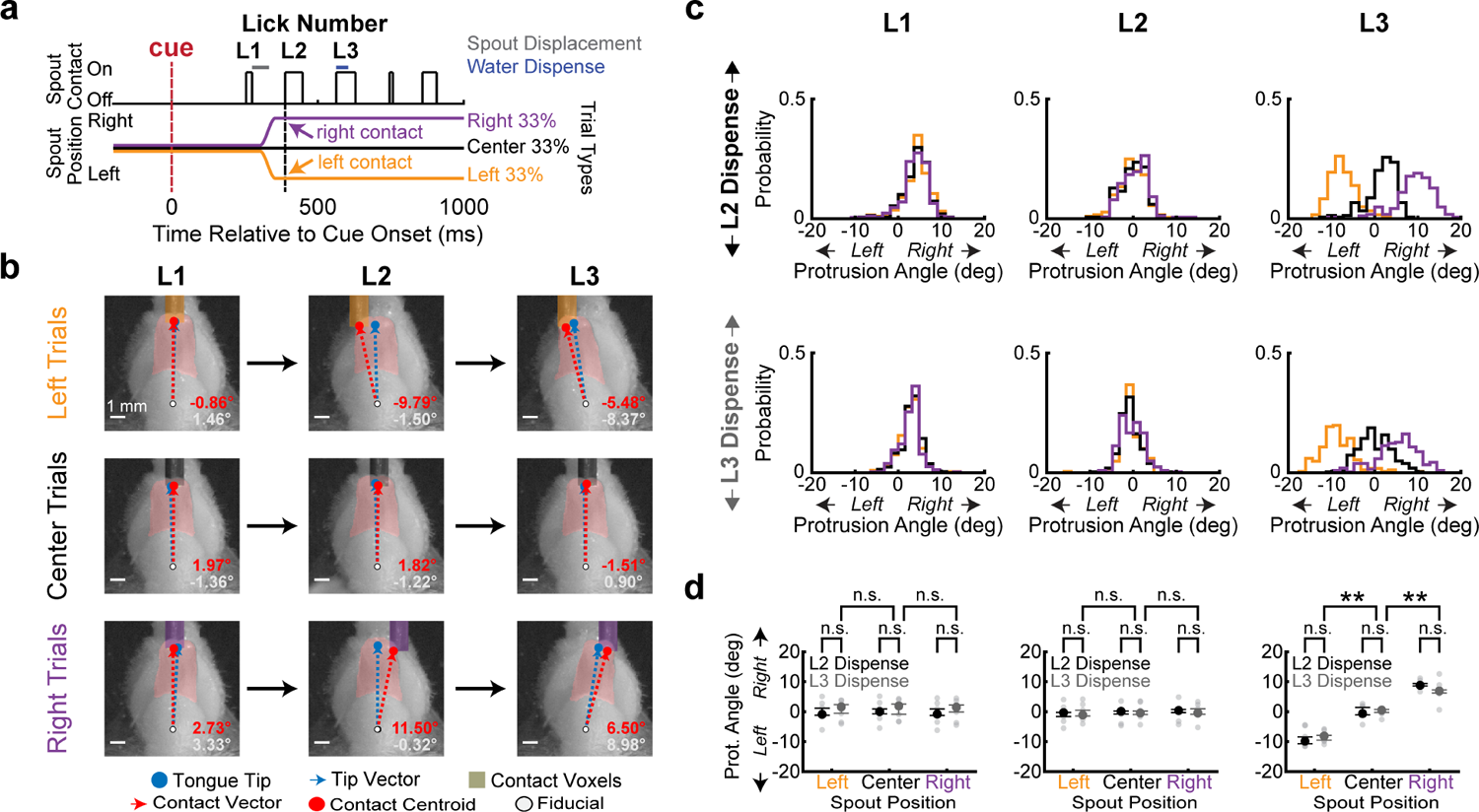
Touch-guided re-aiming does not depend on water dispensation. **a**, Trial structure was the same as in Fig. 1a, except water was dispensed at L3 contact instead of L2 contact. **b**, Bottom view still frames at L1-L3 contact onset across trial types for dispense on L3 trials. Tip site and vector, blue dot and dashed line. Contact site and vector, red dot and dashed line. Contact angle and protrusion angle, red and white values. **c**, L1-L3 protrusion angle distributions during left (gold), center (black) and right (purple) trials from a session where water was dispensed on L2 (top) or L3 (bottom). **d**, L3 protrusion angles for dispense on L2 (black, *n* = 7 mice) and dispense on L3 trials (dark gray, *n* = 7 mice). Gray dots are medians of individual mice. Data in **d** are median ± IQR. **corrected *p* < 0.01, shuffle test; n.s., not significant. Exact statistics are in Supplementary Table 2.

**Extended Data Fig. 2.**
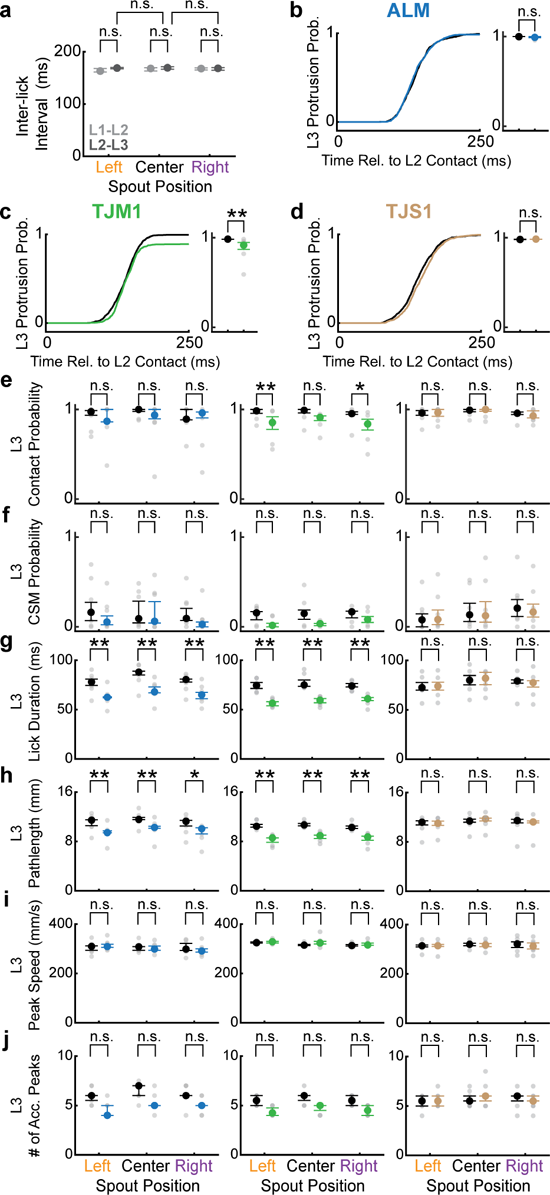
Bilateral photoinactivation of ALM and TJM1, but not TJS1, reduces lick pathlength and duration. **a**, Rhythm of lick bouts are unaffected by spout displacement on L2. Inter-lick intervals (ILI) for L2 to L3 (dark gray) do not significantly differ by condition. ILI for L1 to L2 (light gray), where no spout displacements occurred, provided for comparison. **b-d**, Left, Cumulative probability of L3 protrusion relative to L2 contact onset for ALM (blue, **b**), TJM1 (green, **c**) and TJS1 (brown, **d**) inactivated trials, relative to no laser trials for the same mice (black). Right, L3 protrusion probability for inactivation and no laser trials. **e-j**, Effect of bilateral ALM, TJM1 or TJS1 inactivation on L3 lick kinematics. L3 contact probability (**e**), corrective submovement (CSM) probability (**f**), lick duration (**g**), lick pathlength (**h**), lick peak speed (**i**) and number of acceleration peaks (**j**) for left, center and right trials. Colors for each panel are the same as in **b-d**. Note that ALM and TJM1, but not TJS1, inactivation impair lick duration and pathlength. Data in **e-j** are median ± IQR. *corrected *p* < 0.05, **corrected *p* < 0.01, shuffle test; n.s., not significant. Exact statistics are in Supplementary Tables 1, 3 and 4.

**Extended Data Fig. 3.**
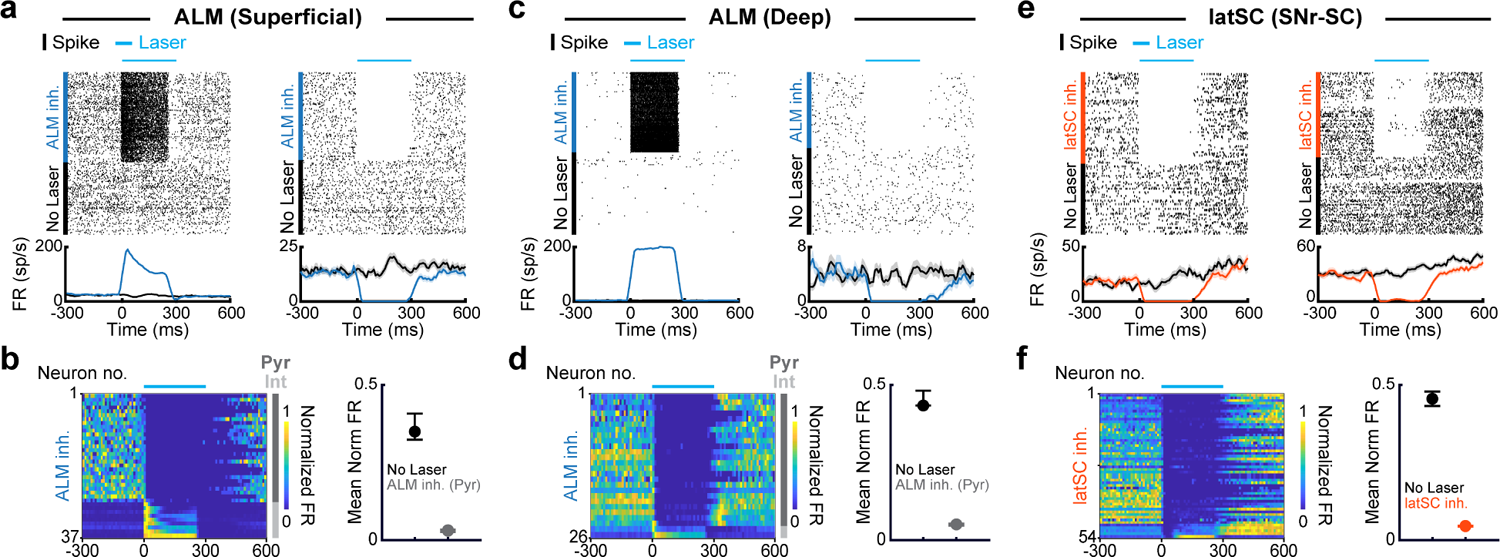
Electrophysiological validation of photoinhibition of the cortex and latSC. **a**, Spike rasters (top row) and rate histograms (bottom row) for an example putative interneuron (left) and pyramidal neuron (right) recorded in the superficial layers of ALM during photovalidation experiments in VGAT-CHR2-EYFP mice. Data are aligned to trial onset and sorted by laser on (dark blue) or off (black) trials. Cyan bar above the raster indicates when the laser was on. Laser power, waveform and duration were the same as in behavioral experiments. **b**, Left, Normalized mean firing rates of superficial layer ALM neurons (300 - 800 μm below dura surface, Methods) recorded during photoinactivation. Dark gray and light gray indicate putative pyramidal and interneurons, respectively. Right, Mean normalized firing rate of putative ALM pyramidal neurons during no laser (black) and laser on (dark gray) trials. **c-d**, Same as **a-b**, but for an example putative interneuron (left) and pyramidal neuron (right) recorded in the deep layers of ALM (800 - 1200 μm below dura surface, Methods). **e-f**, Same as **a-b**, but for two example neurons recorded in latSC during axonal stimulation of the SNr-SC pathway in Vgat-cre mice (orange).

**Extended Data Fig. 4.**
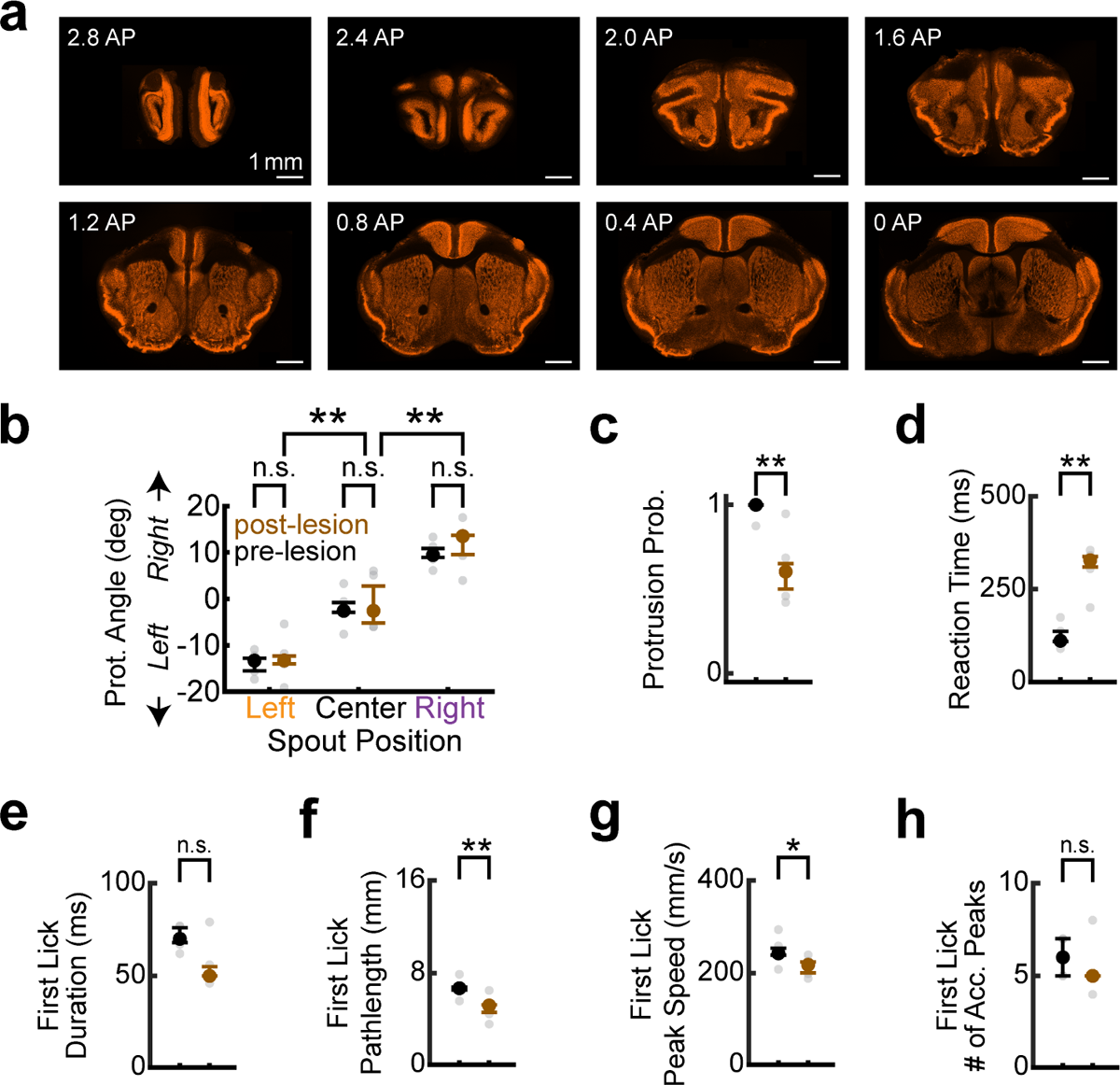
Lesions of all three lingual cortical areas at once impair lick kinematics while preserving touch-guided re-aiming. **a**, Example histological images from a mouse with ALM, TJM1 and TJS1 lesioned. Slices were stained with anti-NeuN (gold). Scale bar, 1 mm. **b-h**, L3 protrusion angles (**b)**, protrusion probability (**c**), reaction time (**d**), first lick duration (**e**), first lick pathlength (**f**), first lick peak speed (**g**), and first lick number of acceleration peaks (**h**) across mice prior to lesion (black) and post-lesion (brown). Data in **b-h** are median ± IQR, *n* = 5 mice. Exact statistics are in Supplementary Tables 5 and 6.

**Extended Data Fig. 5.**
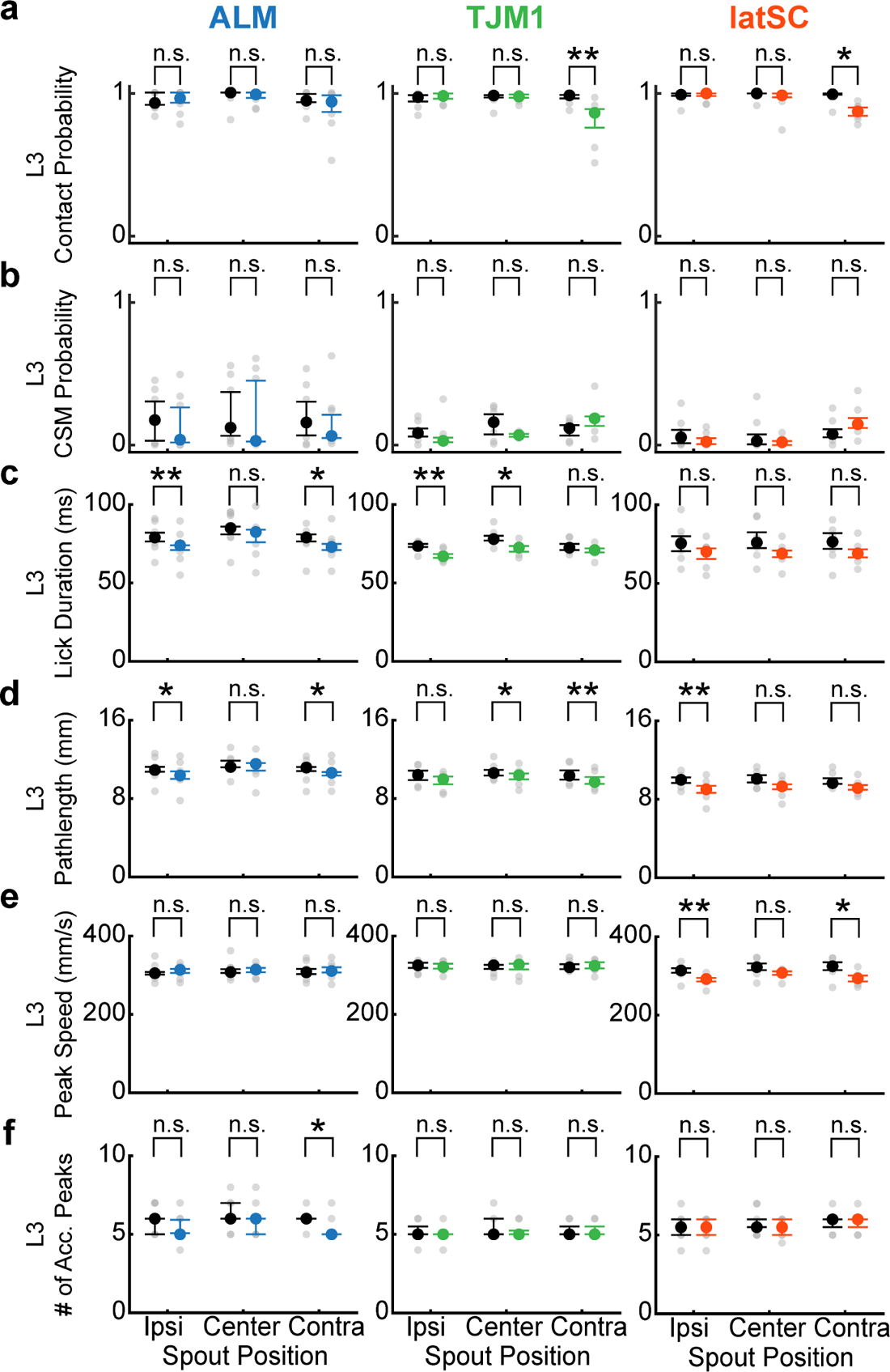
Unilateral photoinactivation of ALM, TJM1 and latSC minimally impairs L3 kinematics. **a-f**, Effect of unilateral ALM, TJM1 or latSC inactivation on L3 lick kinematics for left, center and right trials. **a**, L3 contact probability. **b**, Corrective submovement (CSM) probability. **c**, Lick duration. **d**, Lick pathlength. **e**, Lick peak speed. **f**, Number of acceleration peaks. Data in **a-f** are median ± IQR. Colors in **a-f** correspond to the photoinactivated brain region. Black dots are no laser trials from the same session. \**p* < 0.05, \*\**p* < 0.01, shuffle test; n.s., not significant. Exact statistics are in Supplementary Table 7.

**Extended Data Fig. 6.**
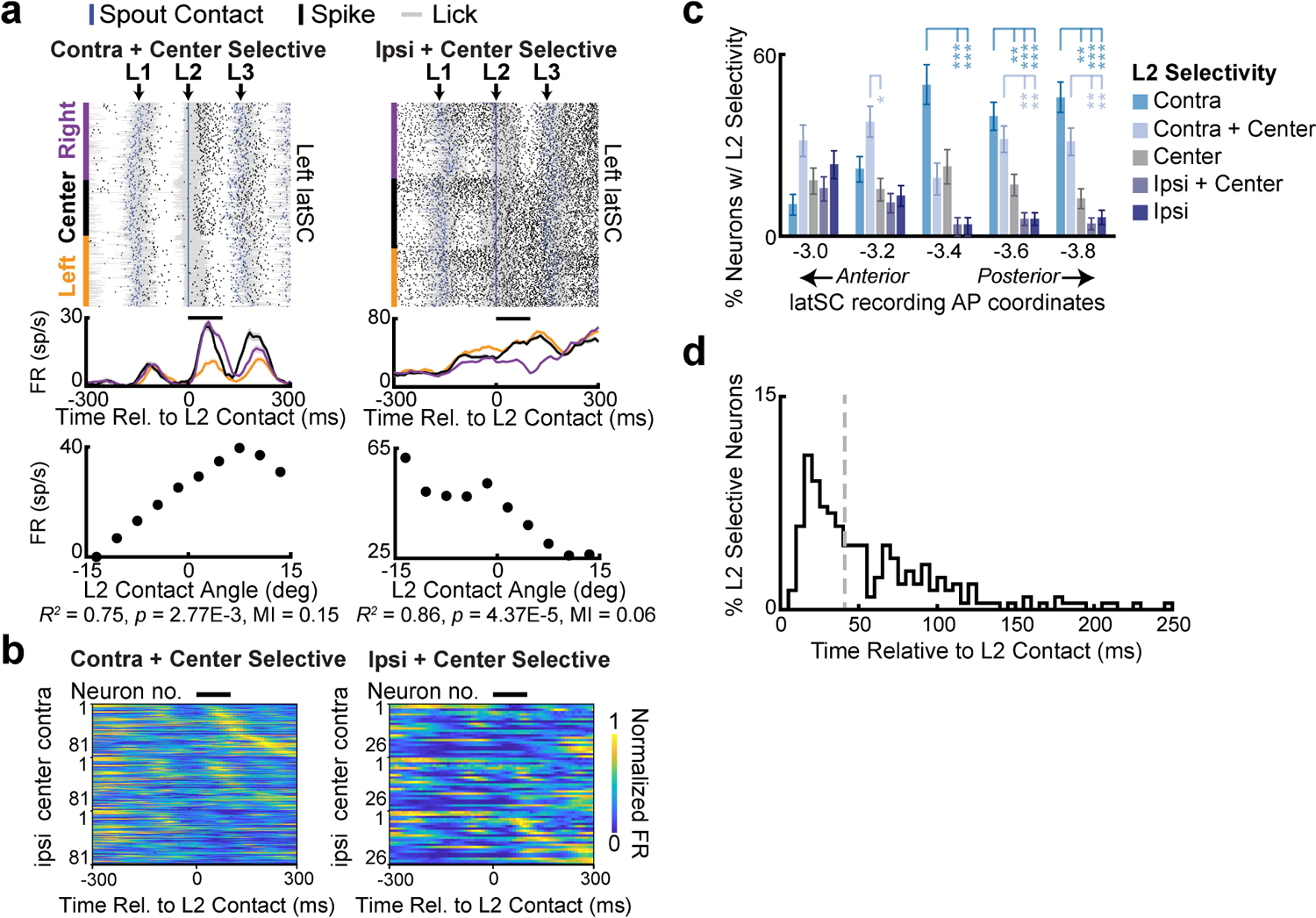
latSC neurons exhibit diverse tuning in response to contact events. **a**, Spike rasters (top row), rate histograms (middle row) and relationship between mean firing rate and L2 contact angle (bottom row) for contralateral + center (left), ipsilateral + center (middle). Data are sorted by condition, then by L2 contact angle within each condition. **b**, Normalized mean firing rates of L2-contact modulated neurons with significant selectivity for contralateral + center (left, *n* = 81/279) and ipsilateral + center (right, *n* = 81/279) contacts on L2. **c**, Percentage of neurons with L2 contralateral, contralateral + center, center, ipsilateral + center, and ipsilateral selectivity as a function of binned recording site (sites are ±: −3.0 (*n* = 38 neurons), −3.2 (*n* = 45), −3.4 (n = 26), −3.6 (*n* = 53), −3.8 (*n* = 48) mm AP. Colors as in **Fig 3c**. Note that there was a significantly larger percentage of neurons with contralateral selectivity in posterior regions of the latSC. *corrected *p* < 0.05, **corrected *p* < 0.01, ***corrected *p* < 0.001, shuffle test. **d**, latSC single neuron divergence latencies for left, center and right trials relative to L2 contact for L2 selective neurons. Gray dashed line, median (41 ms). Note that many single neurons diverged prior to the median, in the 10 - 15 ms range, and that the mode of the distribution was 16 ms.

**Extended Data Fig. 7.**
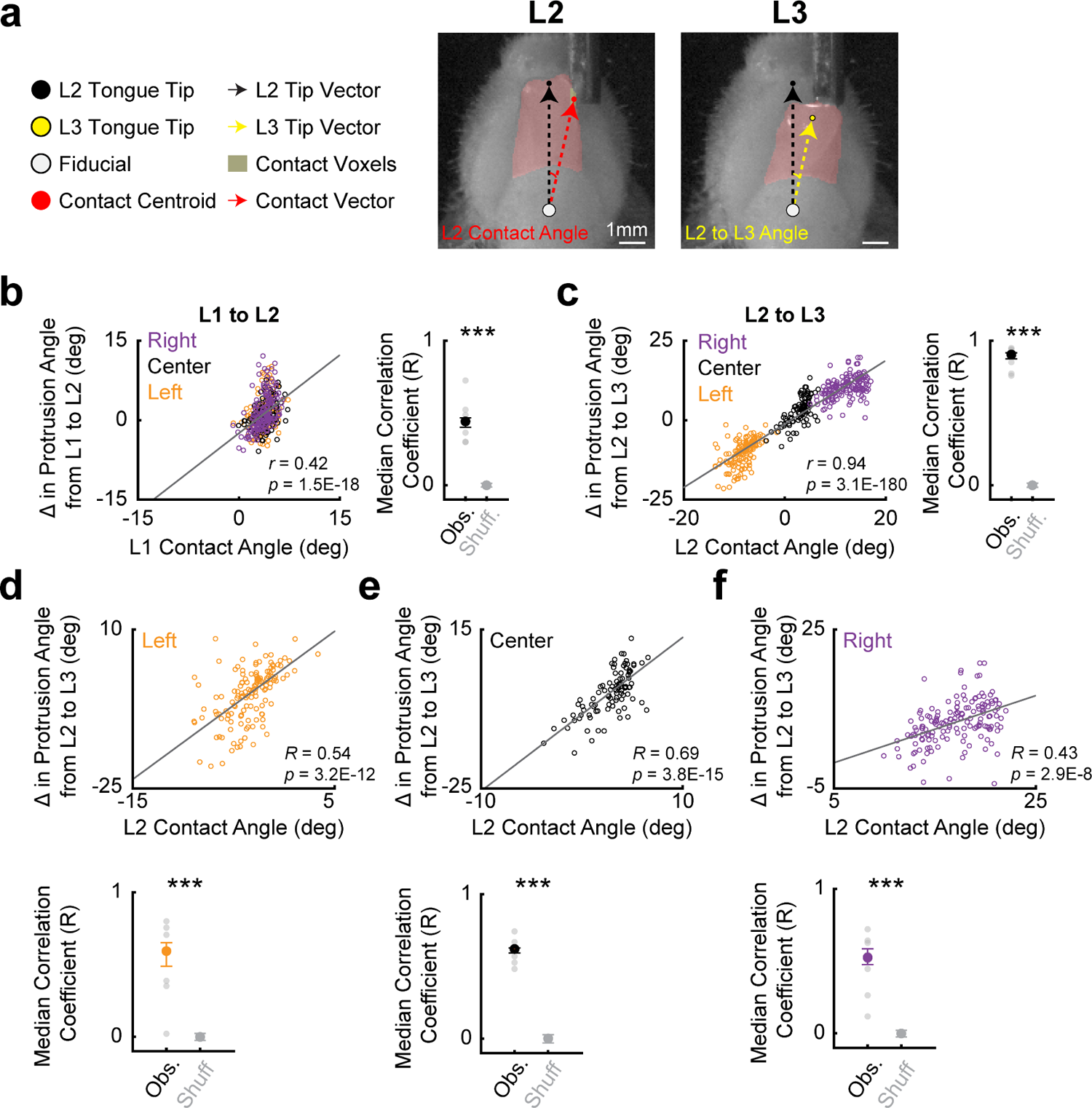
Location of contact events on the tongue predicts next lick re-aiming. **a**, Example still frames of L2 and L3. Contact angle for each lick was defined as the angle between the tip vector (pointing from fiducial to tongue tip) and the contact vector (pointing from fiducial to contact centroid). Protrusion angle change was defined as the angle between the tip vectors at time of spout contact for one lick and the next. **b**, Example data from a single session showing that L1 contact angle (which varied slightly even though the spout was always centered) was significantly correlated with the change in protrusion angle from L1 to L2 across trial types (left). Pearson’s *R* and *p*-value for example data shown in inset. Gray line indicates the line of best fit from linear regression. Right, median correlation coefficient (Pearson’s *R*, black) compared to shuffled data (gray, *n* = 6 mice). Left, center and right displacement trials are yellow, black and purple, respectively. Pearson’s *R* and *p*-value for example data shown in inset. **c**, Same as **b**, but for correlation between L2 contact angle and the change in protrusion angle from L2 to L3 across trial types. Colors as in **b**. **d-f**, Same data as **c** but analyzed separately for each condition: left (**d**), center (**e**), and right (**f**). Top, example data. Bottom, median correlation coefficient compared to shuffled data. \*\*\**p* < 0.001, shuffle test. Exact statistics are in Supplementary Table 9.

**Extended Data Fig. 8.**
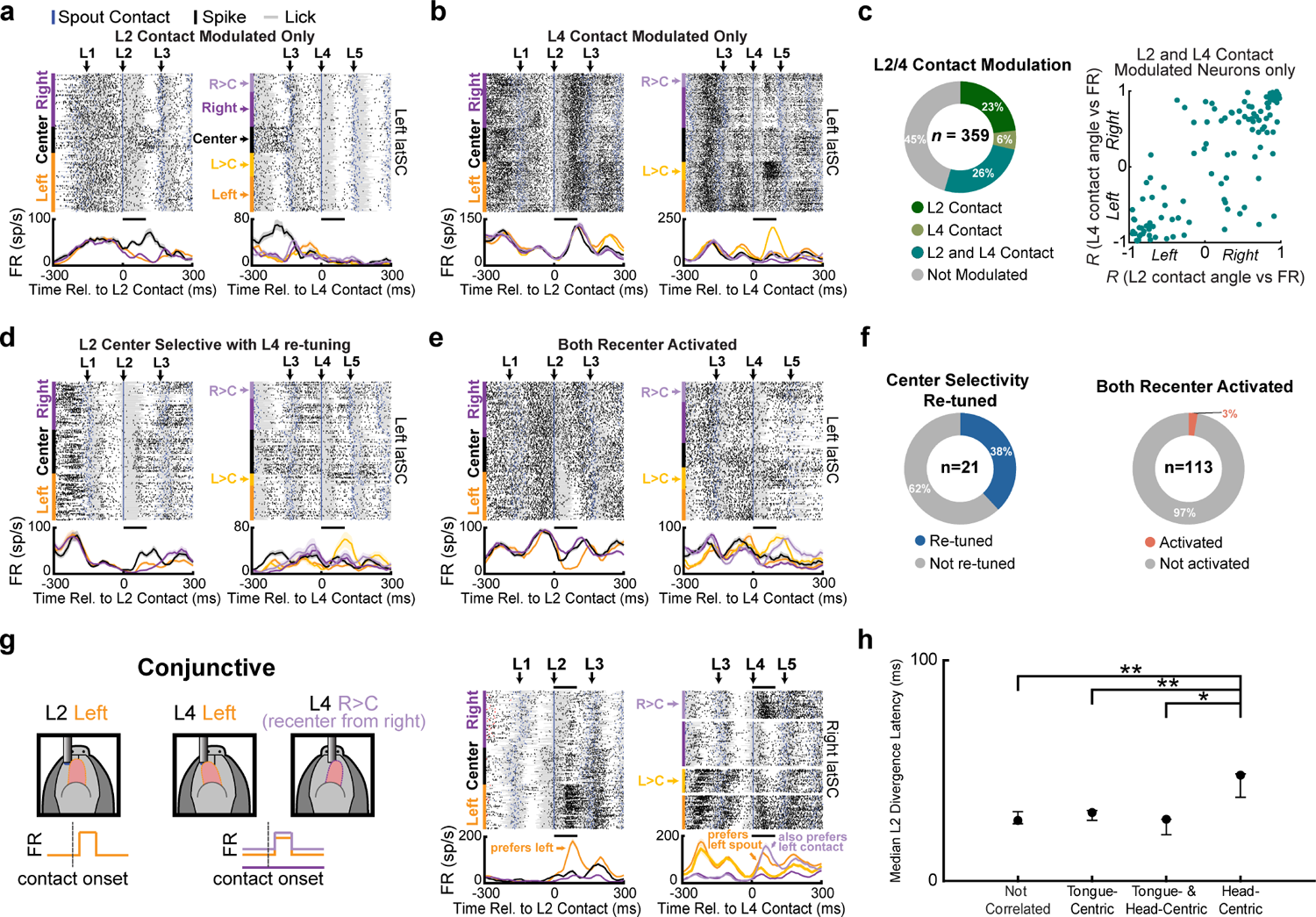
latSC neuronal activity can be selectively tuned to contact events on some licks, but not others. **a-b**, Example latSC neurons tuned to contacts on specific licks. a, A neuron tuned to certain contact locations during L2, but not L4. **b**, A neuron tuned to certain contact locations during L4, but not L2. **c**, Left, Percentage of L2- and L4-modulated neurons tuned to contacts on L2 only (dark green), L4 only (light green), both L2 and L4 (green), or neither L2 or L4 (gray). Right, Pearson’s *R* between L2 contact angle and mean firing rate following L2 contact plotted against Pearson’s *R* for L4 contact angle and mean firing rates following L4 contact. Positive and negative correlation coefficients indicate right or left preference, respectively. Note that the activity of many latSC neurons exhibited similar correlations between firing rate and contact location on L2 and L4. **d**, Example latSC neuron selective for L2 center contacts that was re-tuned to non-center responses on L4 during recentering trials. **e**, Example L4 modulated neuron that was activated by both left and right recentering trials. **f**, Left, Percentage of neurons selective for L2 center contacts that were re-tuned to a non-center contact on L4 (blue). Blue and gray, re-tuned and not re-tuned, respectively. Right, Percentage of L4 modulated neurons that were activated by both left and right recentering trials (orange). **g**, Left, Illustration of a hypothetical conjunctively-tuned neuron. Right, Spike rasters and rate histograms for an example conjunctively-tuned latSC neuron aligned to L2 (left) and L4 contact (right). Colors as in **Fig. 1a**. **h**, Median ± IQR of single neuron divergence latencies between left, center and right conditions for tongue-, head-, and conjunctively-tuned neurons, as well as neurons that were not correlated with any reference frame. Note that tactile responses in neurons with a head-centric reference frame diverged significantly later than those with tongue-centric or conjunctive reference frames. *corrected *p* < 0.05, **corrected *p* < 0.01, hierarchical bootstrap.

**Extended Data Fig. 9.**
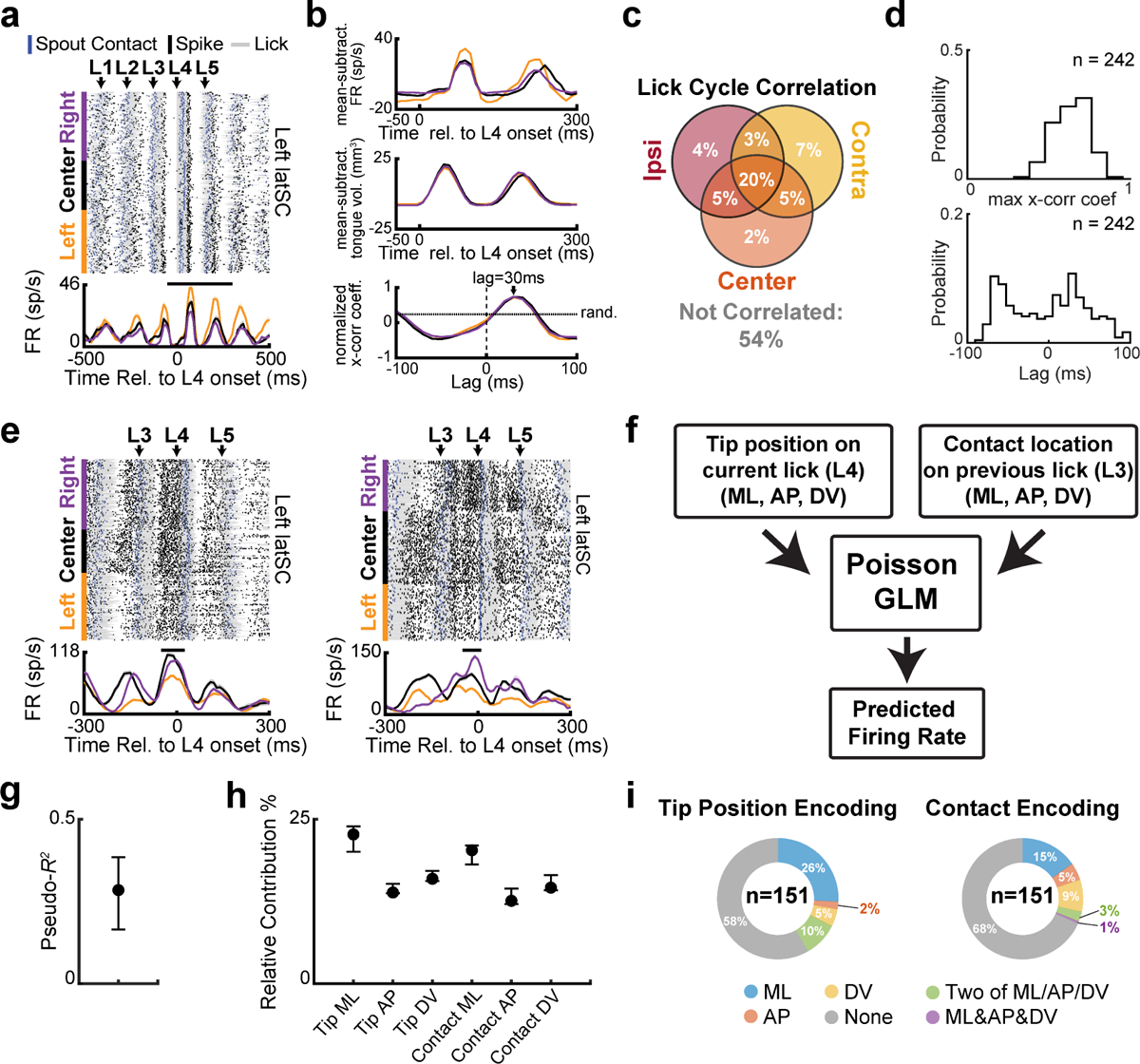
latSC neuronal activity prior to spout contact encodes tongue position. **a**, Example lick cycle correlated latSC. Black bar indicates window used for cross-correlation analysis. b, Top, Mean-subtracted firing rate of the neuron in a. Middle, Mean-subtracted tongue volume. Data in a and b are aligned to L4 protrusion onset. Bottom, Normalized cross-correlation coefficient histogram. Black horizontal line, correlation coefficient from shuffled data. **c**, Percentage of neurons correlated with lick cycle across all combinations of trial types. **d**, Top, Distribution of maximum cross-correlation coefficient across all lick cycle correlated neurons (*n* = 242). Bottom, Distribution of lags for lick cycle correlated neurons. Lag was taken at the time which maximized the cross-correlation coefficient for each neuron. **e**, Two example latSC neurons that are modulated prior to L4. Data are aligned to L4 protrusion onset, and sorted by condition. Within each condition, data are sorted by tip position in the ML plane at L4 protrusion offset. Black bar above the rate histogram indicates the analysis window used for GLMs. **f**, GLM schematic. We sought to test whether tongue position was encoded in pre-contact L4 latSC neuronal activity. However, given the close temporal proximity (∼120 ms) between past (L3) contact and L4 re-aiming, any relationship we might observe between tongue position and latSC neuronal activity may be confounded by past L3 contact location. We included both variables to test whether L4 tongue position was better encoded in latSC pre-contact L4 activity than past L3 contact location. Note that, we are using GLMs as an analysis method to examine encoding of tongue position, not as a model of the latSC, per se. **g**, Mean ± IQR McFadden’s pseudo-*R^2^* across all latSC neurons. pseudo-*R^2^* is a measure of goodness-of-fit for GLMs, with higher values indicating better model fits. Note that pseudo-*R^2^* values of 0.2 - 0.4 represent excellent model fits. **h**, Average relative contribution of each predictor across all latSC neurons. Tip and contact position are separated into their ML, AP or DV components. **i**, Percentage of neurons encoding L4 tip position (left) and previous (L3) contact position. Tip and contact position are broken down into neurons encoding only ML, AP or DV components of position, two of these components, or all three.

**Extended Data Fig. 10.**
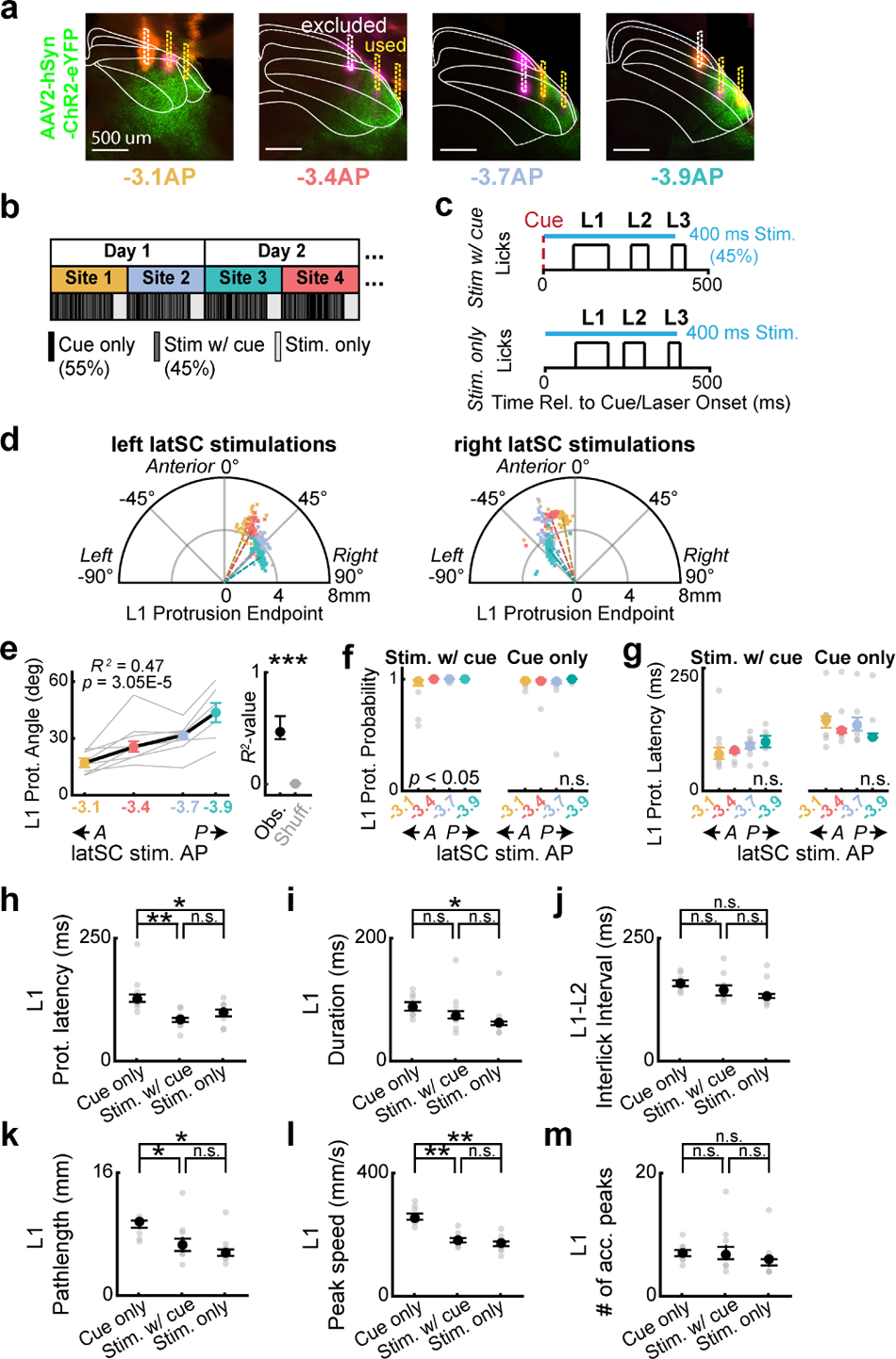
latSC photostimulation reveals the same topographic map when conducted in a cued lick task. **a**, Example histological confirmation of latSC photoactivation sites (−3.1, −3.4, −3.7 and −3.9 mm AP) and AAV2-hSyn-ChR2 expression (green). DiI and DiD tracts corresponding to a single fiber implant site are orange and purple, respectively. Sites were excluded if fiber tracts were outside of the latSC or in regions with little viral expression (white, in contrast to yellow). **b**, Schematic of experimental timeline. Two latSC photostimulation sites were probed per session, with two trial blocks each. In block 1, mice performed ∼120 trials of a cued lick task, where no laser trials (Cue only) were interleaved with latSC photostimulation trials (Stim w/ cue, Methods). In block 2, mice were photostimulated (Stim only) with no task structure (40 trials). This was repeated for each site on each day. **c**, Trial structure for Stim w/ cue (top) and Stim only (bottom) conditions. Red and cyan dashed lines are cue and laser onset, respectively. Cyan bar indicates photostimulation duration. Black pulse trains indicate lick duration for each lick. **d**, Polar plots of L1 protrusion endpoints for an example left (left) and right latSC (right) photostimulation session in the ‘Stim w/ cue’ condition. Colors as in **a**. **e**, latSC-evoked L1 protrusion angles similarly depend on latSC photostimulation site in a cued lick task (*n* = 8 mice, left). Black line and colored dots are median ± IQR across mice for each stimulation site. Gray lines are individual mice. Right, *R^2^* for the observed correlation between photostimulation site and L1 protrusion angle (black) compared to shuffled data (gray) for the ‘Stim w/ cue’ condition. \*\*\**p* < 0.001. **f-g**, L1 protrusion probability (**f**) and protrusion latency (**g**) did not significantly differ in post-hoc tests between latSC stimulation sites for the ‘Stim w/ cue’ and ‘Cue only’ conditions. Kruskal–Wallis test. **h-m**, Lick kinematics during latSC photostimulation for each condition. L1 protrusion latency (**h**), L1 duration (**i**), the L1 - L2 ILI (**j**), L1 pathlength (**k**), peak L1 speed (**l**) and the number of acceleration peaks in L1 (**m**) are shown for all trial types. Note that protrusion latencies were briefer and licks were slightly slower following latSC stimulation than with a cue alone. *corrected *p* < 0.05, **corrected *p* < 0.01, shuffle test; n.s., not significant. Exact statistics are in Supplementary Table 13-14.

## Notes

### Competing Interest Statement

The authors have declared no competing interest.

### Summary of Updates

Extended Data Fig. 1 added, showing that water dispensation was not required for touch-guided re-aiming. Extended Data Fig. 3 added, showing electrophysiological validation of methods used for cortical and latSC photoinhibition. Extended Data Fig. 4 added, showing that touch-guided re-aiming was intact with lesions of all three lingual cortical areas at once. Extended Data Fig. 6d added, showing that many single latSC neurons diverge within 10 - 15 ms relative to contact. Extended Data Fig. 8h added, showing that latSC neurons with contact responses in a head-centric reference frame have longer divergence latencies than neurons with tongue-centric or conjunctive reference frames.

