## Supplementary Tables 1- 14. This file contains Supplementary Tables 1-14, which include statistics for each figure panel. for "A collicular map for touch-guided tongue control"

**Supplementary Table 1. Contact and protrusion angles of L1-L3.** To analyze differences between trials with left/right spout displacements and center trials, shuffle tests were performed and *p*-values were compared against the Bonferroni corrected *p*-value ($p < 0.05/(2\times2) = 1.25\text{E-}2$, two comparisons for each lick number, tests on both tails) to determine significance. Differences between L1-L2 and L2-L3 interlick intervals were also tested with shuffle tests on both tails ($p < 0.05/2 = 2.50\text{E-}2$). $n = 14$ animals.

| | Left | | Center | | Right | | Left vs Center | Right vs Center | Figure panel |
|---|---|---|---|---|---|---|---|---|---|
| | Median | IQR [25% 75%] | Median | IQR [25% 75%] | Median | IQR [25% 75%] | *p*-value | *p*-value | |
| Contact Angle (°) | | | | | | | | | |
| L1 | -0.34 | [-0.75 0.16] | -0.24 | [-0.56 0.15] | 0.12 | [-0.15 0.65] | 4.07E-1 | 1.64E-1 | Fig 1d |
| L2 | -11.93 | [-12.65 -11.42] | 0.49 | [-0.61 1.34] | 13.03 | [12.31 13.81] | <0.001 | <0.001 | |
| L3 | -5.73 | [-6.46 -5.33] | 0.45 | [0.21 0.73] | 6.48 | [5.74 7.40] | <0.001 | <0.001 | |
| Protrusion Angle (°) | | | | | | | | | |
| L1 | 0.80 | [0.19 1.15] | 0.65 | [0.42 1.08] | 0.63 | [0.39 0.95] | 4.58E-1 | 4.25E-1 | Fig 1e |
| L2 | 0.07 | [-0.31 0.35] | -0.18 | [-0.55 0.61] | 0.54 | [0.02 0.80]] | 4.99E-1 | 2.66E-1 | |
| L3 | -9.50 | [-10.05 -9.06] | 0.33 | [-0.02 0.54] | 9.68 | [8.68 10.05] | <0.001 | <0.001 | |
| Interlick Interval from Previous Lick (ms) | | | | | | | | | |
| L2 | 163.25 | [161.25 168.25] | 168.25 | [163.25 169.50] | 168 | [165.25 169.25] | 3.59E-1 | 3.94E-1 | ED Fig 2a |
| L3 | 169 | [167.75 170.25] | 169.5 | [166.75 171.50] | 168.5 | [164.75 170.00] | 4.97E-1 | 3.35E-1 | |
| *p* L2 vs L3 | 2.02E-1 | | 2.47E-1 | | 5.00E-1 | | | | |

**Supplementary Table 2. Protrusion angles of L1-3 with water dispense on L3.** To compare the effects of L2 versus L3 water dispense on protrusion angles, shuffle tests were performed on both tails ($p < 0.05/2 = 2.50E-2$) on both tails to determine significance. To analyze protrusion angle differences between left/right and center, shuffle tests were performed and *p*-values were compared against the Bonferroni corrected *p*-value ($p < 0.05/(2×2) = 1.25E-2$, two comparisons for each lick number, tests on both tails) to determine significance. *n* = 7 animals.

| Protrusion Angle (deg) | Water Dispense | Left | | pre- vs post- | Center | | pre- vs post- | Right | | pre- vs post- | Figure Panel |
|---|---|---|---|---|---|---|---|---|---|---|---|
| | L2 vs L3 | Median | IQR [25% 75%] | *p*-value | Median | IQR [25% 75%] | *p*-value | Median | IQR [25% 75%] | *p*-value | |
| L1 | L2 | -0.89 | [-1.18 1.20] | 2.80E-1 | 0.03 | [-0.69 0.96] | 4.02E-1 | -0.80 | [-1.22 0.92] | 2.33E-1 | ED Fig 1d |
| | L3 | 1.61 | [-0.53 2.40] | | 1.92 | [-0.86 2.57] | | 1.51 | [-0.07 2.25] | | |
| *p* left vs center | | 3.12E-1 | | | | *p* right vs center | | 3.24E-1 | | | |
| L2 | L2 | -0.31 | [-1.62 -0.19] | 4.79E-1 | 0.12 | [-0.77 0.19] | 4.24E-1 | 0.39 | [-0.30 0.76] | 3.89E-1 | ED Fig 1d |
| | L3 | -1.09 | [-1.45 0.53] | | -0.41 | [-0.93 0.20] | | -0.41 | [-0.86 1.01] | | |
| *p* left vs center | | 3.16E-1 | | | | *p* right vs center | | 3.61E-1 | | | |
| L3 | L2 | -9.84158198682830 | -10.75 -8.41 | 2.48E-1 | -0.623929824921518 | -1.06 1.40 | 3.90E-1 | 8.82453184015075 | 8.62 9.39 | 5.20E-2 | ED Fig 1d |
| | L3 | -8.11908086930083 | -9.50 -7.91 | | 0.496328428670781 | -0.24 0.75 | | 6.92083769648174 | 6.27 7.29 | | |
| *p* left vs center | | <0.001 | | | | *p* right vs center | | <0.001 | | | |

**Supplementary Table 3. Kinematics of licks with bilateral photoinactivation.** To compare the effects of inactivation on different kinematic parameters, shuffle tests were performed on both tails ($p < 0.05/2 = 2.50\text{E-}2$) on both tails to determine significance. To analyze L3 protrusion angle differences between left/right and center, shuffle tests were performed and *p*-values were compared against the Bonferroni corrected *p*-value ($p < 0.05/(2\times2) = 1.25\text{E-}2$, two comparisons for each brain region, tests on both tails) to determine significance. $n = 7$ animals, ALM; $n = 6$, TJM1; $n = 6$, TJS1.

| | Laser | Left | | Off vs On | Center | | Off vs On | Right | | Off vs On | Figure Panel |
|---|---|---|---|---|---|---|---|---|---|---|---|
| | Off vs On | Median | IQR [25% 75%] | *p*-value | Median | IQR [25% 75%] | *p*-value | Median | IQR [25% 75%] | *p*-value | |
| L3 Protrusion Angle (deg) | | | | | | | | | | | |
| ALM | Off | -9.31 | [-9.85 -7.40] | 3.25E-1 | 0.28 | [0.19 1.12] | 8.60E-2 | 9.79 | [9.51 10.31] | 3.10E-2 | Fig 1f |
| | On | -8.07 | [-8.67 -7.04] | | -0.14 | [-2.47 0.58] | | 8.70 | [4.20 9.29] | | |
| *p* left vs center | | <0.001 | | | | *p* right vs center | | <0.001 | | | |
| TJM1 | Off | -10.72 | [-12.30-9.12] | 4.53E-1 | -0.40 | [-1.34 -0.02] | 3.82E-1 | 7.98 | [5.90 10.62] | 2.49E-1 | |
| | On | -11.22 | [-11.61 -9.60] | | -2.36 | [-3.48 -0.32] | | 8.60 | [6.40 10.33] | | |
| *p* left vs center | | <0.001 | | | | *p* right vs center | | <0.001 | | | |
| TJS1 | Off | -10.16 | [-10.67 -8.90] | 4.24E-1 | 0.37 | [-0.02 0.61] | 9.4E-2 | 7.28 | [6.74 8.34] | 4.08E-1 | |
| | On | -9.10 | [-9.83 -8.20] | | 0.70 | [0.37 0.90] | | 7.37 | [7.14 8.07] | | |
| *p* left vs center | | <0.001 | | | | *p* right vs center | | <0.001 | | | |
| L3 Contact Probability | | | | | | | | | | | |
| ALM | Off | 0.97 | [0.94 0.99] | 4.49E-1 | 1.00 | [0.98 1.00] | 4.57E-1 | 0.89 | [0.88 1.00] | 4.75E-1 | ED Fig 2e |
| | On | 0.87 | [0.86 1.00] | | 0.94 | [0.90 1.00] | | 0.96 | [0.90 0.97] | | |
| TJM1 | Off | 0.98 | [0.96 1.00] | <0.001 | 0.99 | [0.96 1.00] | 2.80E-2 | 0.95 | [0.95 0.97] | 1.30E-2 | |

| | | | | | | | | | | | |
|---|---|---|---|---|---|---|---|---|---|---|---|
| | On | 0.85 | [0.78 0.92] | | 0.91 | [0.88 0.93] | | 0.84 | [0.77 0.89] | | |
| TJS1 | Off | 0.96 | [0.94 0.99] | 5.79E-1 | 0.99 | [0.98 1.00] | 6.62E-1 | 0.96 | [0.94 0.97] | 4.28E-1 | |
| | On | 0.97 | [0.92 1.00] | | 1.00 | [0.98 1.00] | | 0.93 | [0.91 0.99] | | |
| L3 CSM Probability | | | | | | | | | | | |
| ALM | Off | 0.16 | [0.07 0.27] | 1.90E-1 | 0.09 | [0.04 0.29] | 4.27E-1 | 0.09 | [0.07 0.21] | 6.20E-2 | ED Fig 2f |
| | On | 0.05 | [0.02 0.12] | | 0.06 | [0.04 0.28] | | 0.03 | [0.00 0.05] | | |
| TJM1 | Off | 0.16 | [0.08 0.17] | 9.6E-2 | 0.15 | [0.08 0.19] | 9.2E-2 | 0.17 | [0.10 0.17] | 2.73E-1 | |
| | On | 0.02 | [0.00 0.04] | | 0.03 | [0.01 0.04] | | 0.08 | [0.05 0.11] | | |
| TJS1 | Off | 0.08 | [0.00 0.14] | 2.34E-1 | 0.13 | [0.06 0.26] | 4.84E-1 | 0.21 | [0.11 0.30] | 3.66E-1 | |
| | On | 0.08 | [0.03 0.19] | | 0.12 | [0.05 0.28] | | 0.16 | [0.11 0.25] | | |
| L3 Lick Duration (ms) | | | | | | | | | | | |
| ALM | Off | 78.00 | [74.50 81.00] | <0.001 | 88.00 | [85.00 89.00] | <0.001 | 80.50 | [78.00 81.00] | <0.001 | ED Fig 2g |
| | On | 62.50 | [62.00 63.00] | | 68.00 | [67.00 73.00] | | 65.00 | [61.00 67.50] | | |
| TJM1 | Off | 74.50 | [71.50 77.00] | <0.001 | 75.00 | [73.50 79.88] | <0.001 | 74.00 | [73.00 76.50] | <0.001 | |
| | On | 56.50 | [54.25 58.00] | | 59.50 | [57.00 61.25] | | 61.00 | [59.00 62.13] | | |
| TJS1 | Off | 72.50 | [70.50 77.50] | 4.22E-1 | 80.00 | [72.50 85.00] | 2.29E-1 | 79.25 | [77.50 80.50] | 3.76E-1 | |
| | On | 74.00 | [70.50 78.75] | | 82.00 | [75.75 88.00] | | 77.50 | [73.25 79.50] | | |
| L3 Pathlength (mm) | | | | | | | | | | | |
| ALM | Off | 11.45 | [10.57 11.50] | 8.00E-3 | 11.56 | [11.51 11.89] | 7.00E-3 | 11.27 | [10.50 11.41] | 5.00E-3 | ED Fig 2h |
| | On | 9.43 | [9.40 9.71] | | 10.26 | [10.12 10.46] | | 10.06 | [9.22 10.21] | | |

| | | | | | | | | | | |
|---|---|---|---|---|---|---|---|---|---|---|
| TJM1 | Off | 10.46 | [10.30 10.77] | <0.001 | 10.67 | [10.53 10.93] | <0.001 | 10.26 | [10.11 10.49] | <0.001 | |
| | On | 8.57 | [7.87 8.58] | | 8.93 | [8.46 9.02] | | 8.73 | [8.21 8.86] | | |
| TJS1 | Off | 11.18 | [10.73 11.42] | 4.98E-1 | 11.38 | [11.22 11.72] | 1.28E-1 | 11.46 | [11.09 11.69] | 3.83E-1 | |
| | On | 11.01 | [10.69 11.39] | | 11.73 | [11.37 11.87] | | 11.23 | [11.12 11.39] | | |
| L3 Peak Speed (mm/s) | | | | | | | | | | | |
| ALM | Off | 309.83 | [293.77 312.12] | 2.31E-1 | 308.24 | [293.94 308.54] | 4.54E-1 | 298.62 | [292.86 321.97] | 2.37E-1 | ED Fig 2i |
| | On | 309.07 | [306.09 318.66] | | 298.96 | [293.92 311.06] | | 290.89 | [286.36 299.92] | | |
| TJM1 | Off | 324.56 | [323.60 327.46] | 4.21E-1 | 314.74 | [313.41 318.35] | 1.38E-1 | 313.15 | [311.48 317.07] | 2.84E-1 | |
| | On | 328.68 | [323.60 330.37] | | 323.76 | [319.94 330.71] | | 315.32 | [314.02 323.71] | | |
| TJS1 | Off | 314.31 | [309.81 317.40] | 4.23E-1 | 320.03 | [314.48 325.19 ] | 3.38E-1 | 321.34 | [308.19 330.62 ] | 2.48E-1 | |
| | On | 314.16 | [311.45 319.88 ] | | 317.49 | [311.21 324.19 ] | | 312.36 | [300.90 326.39 ] | | |
| L3 Number of Acceleration Peaks | | | | | | | | | | | |
| ALM | Off | 6.00 | [5.50 6.00] | 7.40E-2 | 7.00 | [6.00 7.00] | 2.90E-2 | 6.00 | [6.00 6.00] | 9.40E-2 | ED Fig 2j |
| | On | 4.00 | [4.00 5.00] | | 5.00 | [5.00 5.00] | | 5.00 | [5.00 5.00] | | |
| TJM1 | Off | 5.50 | [5.00 6.00] | 1.25E-1 | 6.00 | [5.50 6.00] | 5.00E-2 | 5.50 | [5.00 6.00] | 7.30E-2 | |
| | On | 4.25 | [4.00 4.75] | | 5.00 | [4.50 5.00] | | 4.50 | [4.00 5.00] | | |
| TJS1 | Off | 5.50 | [5.00 6.00] | 7.79E-1 | 5.50 | [5.50 6.00] | 5.38E-1 | 6.00 | [5.38 6.00] | 6.80E-1 | |
| | On | 5.50 | [5.00 6.00] | | 6.00 | [5.50 6.00] | | 5.50 | [5.50 6.00] | | |

**Supplementary Table 4. L3 protrusion probability with bilateral photoinactivation.** To analyze the differences between laser off and on conditions, shuffle tests were performed on both tails (*p* < 0.05/2 = 2.50E-2 on both tails to determine significance. *n* = 7 animals, ALM; *n* = 6, TJM1; *n* = 6, TJS1; *n* = 6, latSC.

| | L3 Protrusion Probability | | | | Off vs On | Figure Panel |
|---|---|---|---|---|---|---|
| | Laser Off | | Laser ON | | | |
| | Median | IQR [25% 75%] | Median | IQR [25% 75%] | *p*-value | |
| ALM | 1.00 | [1.00 1.00] | 0.99 | [0.99 1.00] | 5.30E-1 | ED Fig 2b |
| TJM1 | 1.00 | [1.00 1.00] | 0.93 | [0.89 0.96] | 4.00E-3 | ED Fig 2c |
| TJS1 | 1.00 | [1.00 1.00] | 1.00 | [1.00 1.00] | 6.00E-1 | ED Fig 2d |
| SC | 1.00 | [1.00 1.00] | 0.10 | [0.00 0.22] | <0.001 | Fig 2b |

**Supplementary Table 5. L3 protrusion angles with cortical lesions.** To compare the effects of lesions on protrusion angles, shuffle tests were performed on both tails ($p < 0.05/2 = 2.50E\text{-}2$) on both tails to determine significance. *n* = 5 animals.

| | Lesion | Left | | pre- vs post- | Center | | pre- vs post- | Right | | pre- vs post- | Figure Panel |
|---|---|---|---|---|---|---|---|---|---|---|---|
| | pre- vs post- | Median | IQR [25% 75%] | *p*-value | Median | IQR [25% 75%] | *p*-value | Median | IQR [25% 75%] | *p*-value | |
| L3 Protrusion Angle (deg) | pre | -13.31 | [-15.51 -12.79] | 4.82E-1 | -2.57 | [-2.88 -0.78] | 4.63E-1 | 9.45 | [8.93 10.90] | 2.03E-1 | ED Fig 4b |
| | post | -13.28 | [-13.98 -12.39] | | -2.58 | [-5.22 2.79] | | 11.53 | [9.56 13.69] | | |
| *p* left vs center (post-lesion) | | <0.001 | | | | *p* right vs center | | <0.001 | | | |

**Supplementary Table 6. Lick kinematics with cortical lesions.** To compare the effects of lesions on different kinematics, shuffle tests were performed on both tails ($p < 0.05/2 = 2.50E\text{-}2$) on both tails to determine significance. $n = 5$ animals.

| | | Median | IQR [25% 75%] | *p*-value | Figure Panel |
|---|---|---|---|---|---|
| Protrusion Probability | Pre-lesion | 1.00 | [1.00 1.00] | <0.001 | ED Fig 4c |
| | Post-lesion | 0.60 | [0.50 0.65] | | |
| Reaction Time (ms) | Pre-lesion | 111.00 | [109.00 136.00] | <0.001 | ED Fig 4d |
| | Post-lesion | 328.00 | [310.00 338.00] | | |
| L1 Duration (ms) | Pre-lesion | 70.00 | [68.00 76.00] | 6.90E-2 | ED Fig 4e |
| | Post-lesion | 50.00 | [50.00 55.00] | | |
| L1 Pathlength (mm) | Pre-lesion | 6.70 | [6.54 6.81] | <0.001 | ED Fig 4f |
| | Post-lesion | 5.21 | [4.63 5.28] | | |
| L1 Peak Speed (mm/s) | Pre-lesion | 242.63 | [238.63 253.93] | 1.00E-3 | ED Fig 4g |
| | Post-lesion | 217.92 | [200.66 224.45] | | |
| L1 Number of Acceleration Peaks | Pre-lesion | 6.00 | [5.00 7.00] | 3.73E-1 | ED Fig 4h |
| | Post-lesion | 5.00 | [5.00 5.00] | | |

**Supplementary Table 7. Kinematics of licks with unilateral photoinactivation.** To analyze the effects of inactivation on different kinematic parameters, shuffle tests were performed on both tails ($p < 0.05/2 = 2.50E-2$) on both tails to determine significance. $n = 7$ animals, ALM; $n = 6$, TJM1; $n = 6$, latSC.

| | Laser | Ipsilateral | | Off vs On | Center | | Off vs On | Contralateral | | Off vs On | Figure Panel |
|---|---|---|---|---|---|---|---|---|---|---|---|
| | Off vs On | Median | IQR [25% 75%] | *p*-value | Median | IQR [25% 75%] | *p*-value | Median | IQR [25% 75%] | *p*-value | |
| *L3 Protrusion Angle (deg)* | | | | | | | | | | | |
| ALM | Off | -8.18 | [-8.56 - 7.66] | 8.00E-3 | 1.13 | [0.80 1.35] | <0.001 | 9.03 | [8.27 10.21] | <0.001 | Fig 2e |
| | On | -11.76 | [-12.11 - 9.18] | | -3.75 | [-4.04 - 3.12] | | 3.84 | [3.69 4.13] | | |
| TJM1 | Off | -10.56 | [-11.60 - 9.72] | <0.001 | 0.11 | [-0.99 0.41] | <0.001 | 8.90 | [8.25 10.84] | <0.001 | |
| | On | -15.22 | [-16.13 - 14.60] | | -8.46 | [-10.07 - 7.16] | | -0.12 | [-0.68 2.18] | | |
| latSC | Off | -8.21 | [-9.06 - 7.32] | 1.00E-3 | -0.54 | [-0.76 - 0.36] | <0.001 | 6.82 | [6.54 7.94] | <0.001 | |
| | On | -10.20 | [-11.74 - 8.70] | | -3.83 | [-4.35 - 3.73] | | 0.20 | [-1.39 0.31] | | |
| *L3 Contact Probability* | | | | | | | | | | | |
| ALM | Off | 0.93 | [0.91 1.00] | 5.61E-1 | 1.00 | [1.00 1.00] | 4.99E-1 | 0.94 | [0.93 0.99] | 2.38E-1 | ED Fig 5a |
| | On | 0.96 | [0.92 1.00] | | 0.99 | [0.96 1.00] | | 0.94 | [0.86 0.98] | | |
| TJM1 | Off | 0.97 | [0.95 0.99] | 1.91E-1 | 0.99 | [0.97 0.99] | 4.11E-1 | 0.99 | [0.97 0.99] | <0.001 | |
| | On | 0.98 | [0.96 1.00] | | 0.98 | [0.97 0.99] | | 0.86 | [0.77 0.89] | | |
| latSC | Off | 0.99 | [0.99 1.00] | 7.83E-1 | 1.00 | [1.00 1.00] | 2.94E-1 | 0.99 | [0.99 1.00] | 2.20E-2 | |
| | On | 1.00 | [0.98 1.00] | | 0.99 | [0.97 1.00] | | 0.87 | [0.84 0.90] | | |
| *L3 CSM Probability* | | | | | | | | | | | |

| | | | | | | | | | | | |
|---|---|---|---|---|---|---|---|---|---|---|---|
| ALM | Off | 0.17 | [0.02 0.29] | 2.94E-1 | 0.12 | [0.06 0.36] | 3.50E-1 | 0.16 | [0.05 0.30] | 2.90E-1 | ED Fig 5b |
| | On | 0.04 | [0.02 0.25] | | 0.03 | [0.03 0.43] | | 0.06 | [0.04 0.21] | | |
| TJM1 | Off | 0.08 | [0.06 0.11] | 7.90E-2 | 0.16 | [0.08 0.22] | 1.76E-1 | 0.12 | [0.07 0.14] | 1.3E-1 | |
| | On | 0.03 | [0.02 0.05] | | 0.07 | [0.06 0.08] | | 0.19 | [0.14 0.20] | | |
| latSC | Off | 0.05 | [0.02 0.10] | 1.97E-1 | 0.03 | [4.70E-3 0.08] | 3.73E-1 | 0.08 | [0.05 0.12] | 1.43E-1 | |
| | On | 0.02 | [5.90E-3 0.04] | | 0.02 | [0.00 0.03] | | 0.15 | [0.12 0.19] | | |
| *L3 Lick Duration (ms)* | | | | | | | | | | | |
| ALM | Off | 79.00 | [76.00 82.00] | <0.001 | 85.00 | [80.50 86.00] | 3.50E-2 | 79.00 | [76.00 80.00] | 1.10E-2 | ED Fig 5c |
| | On | 74.00 | [70.50 74.00] | | 82.50 | [76.00 84.00] | | 73.00 | [71.00 75.00] | | |
| TJM1 | Off | 73.50 | [73.00 75.00] | 4.00E-3 | 78.00 | [77.00 80.00] | 5.00E-3 | 72.50 | [71.00 75.00] | 2.15E-1 | |
| | On | 67.00 | [66.00 68.5] | | 72.50 | [70.00 73.50] | | 71.00 | [69.00 72.00] | | |
| latSC | Off | 75.50 | [70.50 80.00] | 7.50E-2 | 76.00 | [72.50 82.50] | 3.20E-2 | 76.50 | [72.00 82.00] | 5.90E-2 | |
| | On | 70.25 | [65.50 72.25] | | 69.00 | [66.75 71.00] | | 69.00 | [66.50 71.75] | | |
| *L3 Pathlength (mm)* | | | | | | | | | | | |
| ALM | Off | 10.91 | [10.72 11.24] | 1.00E-2 | 11.22 | [11.20 11.87] | 2.96E-1 | 11.17 | [10.80 11.22] | 2.10E-2 | ED Fig 5d |
| | On | 10.37 | [10.02 10.78] | | 11.54 | [10.84 11.62] | | 10.63 | [10.34 10.71] | | |
| TJM1 | Off | 10.42 | [9.88 10.84] | 3.70E-2 | 10.61 | [10.34 10.91] | 7.00E-3 | 10.33 | [9.93 10.88] | <0.001 | |
| | On | 9.98 | [8.45 10.25] | | 10.38 | [9.96 10.57] | | 9.69 | [9.50 10.21] | | |
| latSC | Off | 9.98 | [9.67 10.23] | <0.001 | 10.09 | [9.71 10.46] | 4.40E-2 | 9.64 | [9.56 10.14] | 3.90E-2 | |
| | On | 9.02 | [8.66 9.37] | | 9.32 | [9.02 9.51] | | 9.14 | [8.97 9.46] | | |

| | | | | | | | | | | | |
|---|---|---|---|---|---|---|---|---|---|---|---|
| *L3 Peak Speed (mm/s)* | | | | | | | | | | | |
| ALM | Off | 305.15 | [301.85 308.38] | 7.84E-1 | 308.15 | [305.87 315.70] | 6.08E-1 | 307.56 | [302.00 316.36] | 6.87E-1 | ED Fig 5e |
| | On | 313.90 | [305.84 315.96] | | 314.38 | [307.87 318.58] | | 310.86 | [306.81 320.16] | | |
| TJM1 | Off | 325.32 | [318.84 331.66] | 3.69E-1 | 325.48 | [315.95 326.67] | 5.98E-1 | 320.26 | [315.55 328.12] | 6.25E-1 | |
| | On | 321.12 | [316.80 329.54] | | 327.28 | [314.69 329.35] | | 324.20 | [317.18 332.79] | | |
| latSC | Off | 313.86 | [308.15 319.73] | <0.001 | 322.40 | [315.01 332.08] | 4.00E-1 | 324.79 | [315.07 334.65] | 1.10E-2 | |
| | On | 292.30 | [285.86 295.04] | | 307.86 | [302.60 311.87] | | 293.83 | [286.05 301.29] | | |
| *L3 Number of Acceleration Peaks* | | | | | | | | | | | |
| ALM | Off | 6.00 | [5.00 6.00] | 4.57E-1 | 6.00 | [6.00 7.00] | 5.11E-1 | 6.00 | [6.00 6.00] | 2.16E-1 | ED Fig 5f |
| | On | 5.00 | [5.00 6.00] | | 6.00 | [5.00 6.00] | | 5.00 | [5.00 5.00] | | |
| TJM1 | Off | 5.00 | [5.00 5.50] | 5.24E-1 | 5.00 | [5.00 6.00] | 3.70E-1 | 5.00 | [5.00 5.50] | 6.63E-1 | |
| | On | 5.00 | [5.00 5.00] | | 5.00 | [5.00 5.25] | | 5.00 | [5.00 5.50] | | |
| latSC | Off | 5.50 | [5.00 6.00] | 6.34E-1 | 5.50 | [5.50 6.00] | 4.93E-1 | 6.00 | [5.50 6.00] | 7.98E-1 | |
| | On | 5.50 | [5.00 6.00] | | 5.50 | [5.00 6.00] | | 6.00 | [5.50 6.00] | | |

**Supplementary Table 8. Ipsilateral (C*ipsi*) and contralateral (C*contra*) correction magnitude with unilateral photoinactivation.** For comparisons between laser off and on conditions, shuffle tests were performed on both tails ($p < 0.05/2 = 2.50E-2$). To compare between inactivation conditions (a-c are Cipsi during ALM, TJM1, and latSC inactivation, and d-f are Ccontra for the same inactivations), shuffle tests were performed and p-values were examined against the Bonferroni corrected *p*-value ($p < 0.05/(3×2) = 8.30E-3$, three pairs of comparisons on ipsilateral or contralateral sides, tests on both tails) to determine significance. *n* = 7 animals, ALM; *n* = 6, TJM1; *n* = 6, latSC.

| | Laser | C*ipsi* (deg) | | Off vs On | C*contra* (deg) | | Off vs On | Figure Panel |
|---|---|---|---|---|---|---|---|---|
| | Off vs On | Median | IQR [25% 75%] | *p*-value | Median | IQR [25% 75%] | *p*-value | Fig 2f |
| ALM | Off | 8.93 | [7.62 9.57] | 1.03E-1 | 8.15 | [7.61 8.56] | 1.35E-1 | |
| | On | 6.93 (a) | [6.15 7.51] | | 7.60 (d) | [6.56 7.79] | | |
| TJM1 | Off | 10.14 | [9.84 10.66] | <0.001 | 10.36 | [9.88 11.48] | 2.29E-1 | |
| | On | 7.28 (b) | [6.04 7.48] | | 8.52 (e) | [8.31 10.21] | | |
| latSC | Off | 7.55 | [6.97 7.83] | 4.70E-2 | 7.85 | [7.40 8.64] | <0.001 | |
| | On | 5.29 (c) | [4.97 6.74] | | 4.00 (f) | [3.07 4.05] | | |

| *Comparisons across brain regions* | |
|---|---|
| Pairwise Comparison | *p*-value |
| ab (Cipsi ALM vs Cipsi TJM1) | 4.71E-1 |
| ac (Cipsi ALM vs Cipsi latSC) | 2.71E-1 |
| bc (Cipsi TJM1 vs Cipsi latSC) | 2.92E-1 |
| de (Ccontra ALM vs Ccontra TJM1) | 7.60E-2 |
| df (Ccontra ALM vs Ccontra latSC) | <0.001 |
| ef (Ccontra TJM1 vs Ccontra latSC) | <0.001 |

**Supplementary Table 9. Correlations between protrusion angle changes and contact angles on the tongue.** Observed Pearson's R are compared with shuffled values on both tails ($p < 0.05/2 = 2.50E-2$). $n = 8$ animals.

| Licks | Trial Types | Observed R | | Shuffled R | | *p*-value | Figure Panel |
|---|---|---|---|---|---|---|---|
| | | Median | IQR [25% 75%] | Median | IQR [25% 75%] | | |
| L1-L2 | All | 0.44 | [0.40 0.47] | -2.83E-4 | [-1.29E-2 1.25E-2] | <0.001 | ED Fig 7b |
| L2-L3 | All | 0.90 | [0.87 0.92] | -2.41E-4 | [-1.42E-2 1.15E2] | <0.001 | ED Fig 7c |
| L2-L3 | Left | 0.59 | [0.49 0.65] | -1.60E-3 | [-2.58E-2 2.26E-2] | <0.001 | ED Fig 7d |
| L2-L3 | Center | 0.62 | [0.59 0.63] | 1.40E-3 | [-2.87E-2 2.73E-2] | <0.001 | ED Fig 7e |
| L2-L3 | Right | 0.52 | [0.47 0.59] | -1.40E-3 | [-2.55E-2 2.15E-2] | <0.001 | ED Fig 7f |

**Supplementary Table 10. Contact and protrusion angles of L3-L5 in recentering trials.** To analyze differences between recentering trials (L>C & R>C) with center trials, shuffle tests were performed and *p*-values were compared against the Bonferroni corrected *p*-value ($p < 0.05/(2\times2) = 1.25E-2$, two comparisons for each lick number, tests on both tails) to determine significance. $n = 5$ animals.

| | Left-Recenter (L>C) | | Right-Recenter (R>C) | | L>C vs Center | R>C vs Center | Figure Panel |
|---|---|---|---|---|---|---|---|
| | Median | IQR [25% 75%] | Median | IQR [25% 75%] | *p*-value | *p*-value | |
| Contact Angle (°) | | | | | | | |
| L3 | -4.44 | [-6.24 - 4.03] | 6.30 | [6.05 7.22] | <0.001 | <0.001 | Fig 4b |
| L4 | 7.83 | [7.76 8.33] | -5.69 | [-6.49 - 5.60] | <0.001 | <0.001 | |
| L5 | 3.03 | [2.08 3.03] | 0.12 | [-1.68 0.36] | 1.69E-1 | 6.00E-3 | |
| Protrusion Angle (°) | | | | | | | |
| L3 | -9.56 | [-11.11 -9.29] | 8.53 | [7.24 8.94] | <0.001 | <0.001 | Fig 4b |
| L4 | -8.08 | [-8.90 - 7.91] | 7.57 | [6.84 8.28] | <0.001 | <0.001 | |
| L5 | -2.60 | [-2.89 - 1.81] | 2.20 | [-0.34 2.63] | 7.50E-2 | 1.60E-2 | |

**Supplementary Table 11. Contact and protrusion angles of L3-L5 in non-recentering trials.** To analyze differences between recentering trials (L>C & R>C) with non-recentering trials (Left & Right), shuffle tests were performed on both tails ($p < 0.05/2 = 2.50E-2$) to determine significance. $n = 5$ animals.

| | Left | | Center | | Right | | Left vs L>C | Right vs R>C | Figure Panel |
|---|---|---|---|---|---|---|---|---|---|
| | Median | IQR [25% 75%] | Median | IQR [25% 75%] | Median | IQR [25% 75%] | *p*-value | *p*-value | |
| Contact Angle (°) | | | | | | | | | |
| L3 | -5.18 | [-6.13 -4.77] | 0.39 | [0.09 0.99] | 6.82 | [5.77 7.00] | 3.01E-1 | 4.10E-1 | Fig 4b |
| L4 | -4.21 | [-5.83 -3.92] | 0.19 | [-0.13 0.41] | 7.21 | [5.46 7.52] | <0.001 | <0.001 | |
| L5 | -2.94 | [-3.18 -2.63] | 1.42 | [0.80 2.37] | 3.58 | [3.35 5.13] | <0.001 | <0.001 | |
| Protrusion Angle (°) | | | | | | | | | |
| L3 | -10.18 | [-10.44 -9.51] | -0.13 | [-0.71 -0.09] | 9.30 | [8.78 9.95] | 4.94E-1 | 1.41E-1 | Fig 4b |
| L4 | -9.76 | [-10.23 -9.24] | 0.02 | [-1.11 0.44] | 10.30 | [9.08 10.65] | 8.00E-3 | 3.70E-2 | |
| L5 | -11.56 | [-11.86 -11.45] | -0.41 | [-1.24 -0.26] | 11.63 | [11.46 13.94] | <0.001 | <0.001 | |

**Supplementary Table 12. Recording locations of example neurons.** Coordinates are all relative to bregma, recorded with an H6 acute probe from Cambridge Neurotech.

| Figure Panel | Description | AP (mm) | ML (mm) | DV (mm) |
|---|---|---|---|---|
| Fig 3a | Contra-preferring | -3.7 | -1.6 | -2.65 |
| | Center-preferring | -3.1 | -1.2 | -2.7 |
| | Ipsi-preferring | Recording site not tracked for this session | | |
| Fig 4c | Tongue-centric | -3.7 | -1.6 | -2.65 |
| Fig 4d | Head-centric | -3.1 | -1.25 | -2.7 |
| ED Fig 6a | Contra & center-preferring | Recording site not tracked for this session | | |
| | Ipsi & center-preferring | -3.1 | -1.3 | -2.7 |
| ED Fig 8a | L2 contact-modulated only | -3.55 | -1.5 | -2.6 |
| ED Fig 8b | L4 contact-modulated only | -3.7 | -1.6 | -2.65 |
| ED Fig 8d | L2 center-preferring & L4 retuning | -3.4 | -1.5 | -2.7 |
| ED Fig 8e | Both recenter activated | -3.4 | -1.53 | -2.7 |
| ED Fig 8g | Conjunctive | -3.1 | -1.25 | -2.7 |
| ED Fig 9a | Lick cycle correlated | Recording site not tracked for this session | | |
| ED Fig 9e | Left lick encoding | Recording site not tracked for this session | | |
| | Right lick encoding | -3.4 | -1.5 | -2.6 |

**Supplementary Table 13. L1 protrusion kinematics across stimulation sites**. Kruskal-Wallis (K-W) tests were used for comparisons across stimulation sites. Correlations between protrusion angles and sites were compared with shuffles. *n* = 8 animals.

| | -3.1AP | | -3.4AP | | -3.7AP | | -3.9AP | | K-W test | Figure Panel |
|---|---|---|---|---|---|---|---|---|---|---|
| | Median | IQR [25% 75%] | Median | IQR [25% 75%] | Median | IQR [25% 75%] | Median | IQR [25% 75%] | *p*-value | |
| *Stimulation Only* | | | | | | | | | | |
| L1 Protrusion Angle (deg) | 17.61 | [15.19 19.68] | 23.79 | [21.68 25.68] | 30.74 | [29.23 33.89] | 39.27 | [33.33 50.02] | 4.52E-4 | Fig 5d |
| Observed $R^2$ : 0.64 [0.55 0.71] , Shuffled $R^2$ : 1.14E-2 [5.10E-3 2.32E-2], $p < 0.001$ | | | | | | | | | | |
| L1 Protrusion Probability | 0.98 | [0.86 1.00] | 1.00 | [1.00 1.00] | 1.00 | [1.00 1.00] | 1.00 | [1.00 1.00] | 1.32E-1 | Fig 5e |
| Across all sites: 0.98 [0.96 1.00] | | | | | | | | | | Fig 3b |
| L1 Protrusion Latency (ms) | 129.00 | [92.00 181.38] | 84.00 | [75.00 90.50] | 107.25 | [102.50 121.13] | 129.50 | [121.00 132.00] | 2.30E-1 | Fig 5f |
| *Stimulation with Cue* | | | | | | | | | | |
| L1 Protrusion Angle (deg) | 17.87 | [15.11 19.81] | 25.54 | [22.88 28.67] | 31.74 | [30.19 32.95] | 43.76 | [38.86 48.78] | 2.80E-3 | ED Fig 10e |
| Observed $R^2$ : 0.47 [0.40 0.61] , Shuffled $R^2$ : 4.10E-3 [1.50E-3 8.00E-3], $p < 0.001$ | | | | | | | | | | |
| L1 Protrusion Probability | 0.98 (a) | [0.93 0.99] | 1.00 (b) | [1.00 1.00] | 1.00 (c) | [1.00 1.00] | 1.00 (d) | [1.00 1.00] | 1.91E-2[#] | ED Fig 10f |
| L1 Protrusion Latency (ms) | 74.50 | [64.75 88.00] | 82.50 | [77.75 82.50] | 91.75 | [86.00 98.00] | 99.75 | [88.50 112.00] | 1.76E-1 | ED Fig 10g |
| *Cue Only* | | | | | | | | | | |
| L1 Protrusion Probability | 0.99 | [0.97 0.99] | 0.98 | [0.98 1.00] | 0.98 | [0.96 0.99] | 1.00 | [1.00 1.00] | 2.80E-1 | ED Fig 10f |
| L1 Protrusion | 144.52 | [128.25 151.38] | 123.00 | [118.00 126.75] | 134.25 | [122.88 149.13] | 109.50 | [109.00 116.75] | 7.97E-1 | ED Fig |

| Latency (ms) | | | | | | | | | | 10g |
|---|---|---|---|---|---|---|---|---|---|---|

### Tukey’s HSD post-hoc test of the K-W test comparing L1 protrusion probability across sites in the *Stimulation with Cue* condition (see table above). None of the p-values was less than the Bonferroni corrected *p*-value ($p < 0.05/6 = 8.30E-3$)

| Pairwise Comparison | *p*-value |
|---|---|
| ab (-3.1AP vs -3.4AP) | 8.31E-2 |
| ac (-3.1AP vs -3.7AP) | 3.59E-2 |
| ad (-3.1AP vs -3.9AP) | 5.58E-2 |
| bc (-3.4AP vs -3.7AP) | 9.85E-1 |
| bd (-3.4AP vs -3.9AP) | 9.87E-1 |
| cd (-3.7AP vs -3.9AP) | 1.00 |

**Supplementary Table 14. Kinematics of licks with latSC photomicrostimulation compared with cue-evoked licks.** To analyze effects of cue and/or photostimulation, shuffle tests were performed and *p*-values were compared against the Bonferroni corrected *p*-value ($p < 0.05/(3\times2) = 8.30E\text{-}3$, three comparisons across the three conditions, tests on both tails) to determine significance. $n = 8$ animals.

| | Cue Only (a) | | Stim. w/ Cue (b) | | Stimulation Only (c) | | Pairwise comparison | Figure Panel |
|---|---|---|---|---|---|---|---|---|
| | Median | IQR [25% 75%] | Median | IQR [25% 75%] | Median | IQR [25% 75%] | *ab p-value* | |
| | | | | | | | *ac p-value* | |
| | | | | | | | *bc p-value* | |
| L1 Protrusion Latency (ms) | 126.50 | [120.50 135.50] | 84.50 | [79.25 88.0] | 99.25 | [91.00 104.5] | <0.001 | ED Fig 10h |
| | | | | | | | 6.00E-3 | |
| | | | | | | | 7.60E-2 | |
| L1 Duration (ms) | 88.00 | [82.00 96.0] | 73.50 | [69.50 81.13] | 62.50 | [58.50 64.50] | 8.50E-2 | ED Fig 10i |
| | | | | | | | 4.00E-3 | |
| | | | | | | | 9.00E-3 | |
| L1-L2 Inter-lick Interval (ms) | 158.00 | [152.38 164.63] | 144.50 | [133.50 154.00] | 132.00 | [128.00 136.50] | 1.27E-1 | ED Fig 10j |
| | | | | | | | 2.30E-2 | |
| | | | | | | | 5.30E-2 | |
| L1 Pathlength (mm) | 9.61 | [8.83 9.77] | 6.60 | [5.80 7.42] | 5.57 | [5.15 6.00] | 7.00E-3 | ED Fig 10k |
| | | | | | | | 2.00E-3 | |
| | | | | | | | 1.50E-2 | |
| L1 Peak Speed (mm/s) | 253.37 | [248.18 268.01] | 181.36 | [174.44 188.20] | 172.17 | [162.43 177.52] | <0.001 | ED Fig 10l |
| | | | | | | | <0.001 | |
| | | | | | | | 1.00E-2 | |
| L1 # of acceleration Peaks | 7.00 | [6.50 7.50] | 6.75 | [6.00 8.00] | 6.00 | [5.00 6.00] | 5.19E-1 | ED Fig 10m |
| | | | | | | | 4.20E-2 | |
| | | | | | | | 2.20E-2 | |
